# Kif1a and intact microtubules maintain synaptic-vesicle populations at ribbon synapses in zebrafish hair cells

**DOI:** 10.1101/2024.05.20.595037

**Authors:** Sandeep David, Katherine Pinter, Keziah-Khue Nguyen, David S. Lee, Zhengchang Lei, Yuliya Sokolova, Lavinia Sheets, Katie S. Kindt

## Abstract

Sensory hair cells of the inner ear utilize specialized ribbon synapses to transmit sensory stimuli to the central nervous system. This sensory transmission necessitates rapid and sustained neurotransmitter release, which relies on a large pool of synaptic vesicles at the hair-cell presynapse. Work in neurons has shown that kinesin motor proteins traffic synaptic material along microtubules to the presynapse, but how new synaptic material reaches the presynapse in hair cells is not known. We show that the kinesin motor protein Kif1a and an intact microtubule network are necessary to enrich synaptic vesicles at the presynapse in hair cells. We use genetics and pharmacology to disrupt Kif1a function and impair microtubule networks in hair cells of the zebrafish lateral-line system. We find that these manipulations decrease synaptic-vesicle populations at the presynapse in hair cells. Using electron microscopy, along with *in vivo* calcium imaging and electrophysiology, we show that a diminished supply of synaptic vesicles adversely affects ribbon-synapse function. *Kif1a* mutants exhibit dramatic reductions in spontaneous vesicle release and evoked postsynaptic calcium responses. Additionally, we find that *kif1a* mutants exhibit impaired rheotaxis, a behavior reliant on the ability of hair cells in the lateral line to respond to sustained flow stimuli. Overall, our results demonstrate that Kif1a-based microtubule transport is critical to enrich synaptic vesicles at the active zone in hair cells, a process that is vital for proper ribbon-synapse function.

**Key points:**

- Kif1a mRNAs are present in zebrafish hair cells
- Loss of Kif1a disrupts the enrichment of synaptic vesicles at ribbon synapses
- Disruption of microtubules depletes synaptic vesicles at ribbon synapses
- Kif1a mutants have impaired ribbon-synapse and sensory-system function

## Introduction

Sensory hair cells in the inner ear rely on specialized ribbon synapses to encode auditory and vestibular stimuli. During stimulation, ribbon synapses are capable of releasing synaptic vesicles at high rates with exquisite precision over long durations (Nouvian *et al*., 2006; Mukhopadhyay & Pangrsic, 2022). The morphology of the presynaptic active zone (AZ) of ribbon synapses is specifically designed to sustain these release properties. Additionally, hair cells also have efficient mechanisms to recycle synaptic vesicles in order to sustain release (Wichmann & Moser, 2015; Pangrsic & Vogl, 2018). Disrupting presynapse morphology or vesicle recycling disrupts ribbon-synapse function and can lead to hearing and balance impairments (Trapani *et al*., 2009; Jing *et al*., 2013; Kroll *et al*., 2019; Wan *et al*., 2019). While considerable work has focused on the importance of synaptic vesicle recycling, less is known about how new synaptic material is made and transported to the presynaptic AZ in hair cells.

Hair cells convert sensory stimuli into electrical impulses to transmit to the brain. Sensory stimuli deflect cilia-like hair bundles at the apex of the cell, opening mechanoelectrical transduction (MET) channels and depolarizing the cell in a graded manner (Gillespie & Walker, 2001). Cell depolarization activates voltage-gated Ca_v_1.3 channels at the presynaptic AZ at the base of the hair cell (Brandt *et al*., 2003). Calcium influx through Ca_v_1.3 channels leads to synaptic-vesicle fusion and glutamate release onto the afferent neuron. Hair cells exhibit rapid and sustained neurotransmission in part due to a specialized presynaptic structure known as the synaptic ribbon (Moser *et al*., 2006; Matthews & Fuchs, 2010). The synaptic ribbon is an electron-dense structure that anchors synaptic vesicles at the presynaptic AZ, readying them for release. Ribbon synapses have been shown to be indefatigable, with rapid vesicle replenishment allowing for sustained release (Griesinger *et al*., 2005; Holt *et al*., 2004). But how hair cells supply sufficient synaptic vesicles at the presynaptic AZ to sustain synapse function is not fully understood.

Synaptic vesicle recycling, through endocytosis and local recycling of synaptic materials at the presynapse, has been shown to be critical to maintain synapse function in neurons and hair cells (Neef *et al*., 2014; Gallimore *et al*., 2023). But in neurons, numerous studies have also outlined the importance of *de novo* transport of synaptic material from the cell soma to presynaptic terminals (Santos *et al*., 2009; Rizzoli, 2014). In the soma of neurons, new synaptic material is synthesized in the endoplasmic reticulum (ER), and then packaged and sorted in Golgi. This material leaves the Golgi as synaptic cargos that are transported along axons via microtubule highways and kinesin motor proteins. Kinesins transport cargos, such as synaptic-vesicle precursors, towards the plus-end of microtubules. In neurons, the plus-end of microtubules in axonal nerve processes are oriented towards the presynapse; thus kinesin-mediated transport delivers synaptic cargos to the presynapse (Guedes-Dias & Holzbaur, 2019). KIF1A, a member of the kinesin-3 family, has been shown to play a conserved role in transporting synaptic-vesicle precursors to neuronal presynapses across multiple species (Yonekawa *et al*., 1998; Zhao *et al*., 2001; Zhao *et al*., 2001; Pack-Chung *et al*., 2007; Rizalar *et al*., 2021). In neurons axons can be extremely long, and cargos must be transported considerable distances to reach the presynapse. In contrast, hair cells are compact, polarized epithelial cells. In hair cells the Golgi is located on top of the nucleus, and the presynaptic AZ is just below the nucleus (Figure 1D, (Siegel & Brownell, 1986)). Recent work has demonstrated that hair-cell microtubules grow their plus ends from the cell apex, past the Golgi, and toward the presynaptic AZ (Hussain *et al*., 2024; Voorn *et al*., 2024). Additionally, immunostaining and scRNAseq data have shown that Kif1a is present in hair cells (Michanski *et al*., 2019; Sur *et al*., 2023). However, whether Kif1a or a microtubule network is required for proper transport of synaptic-vesicle precursors to the presynaptic AZ in hair cells remains unknown.

**Figure 1.**
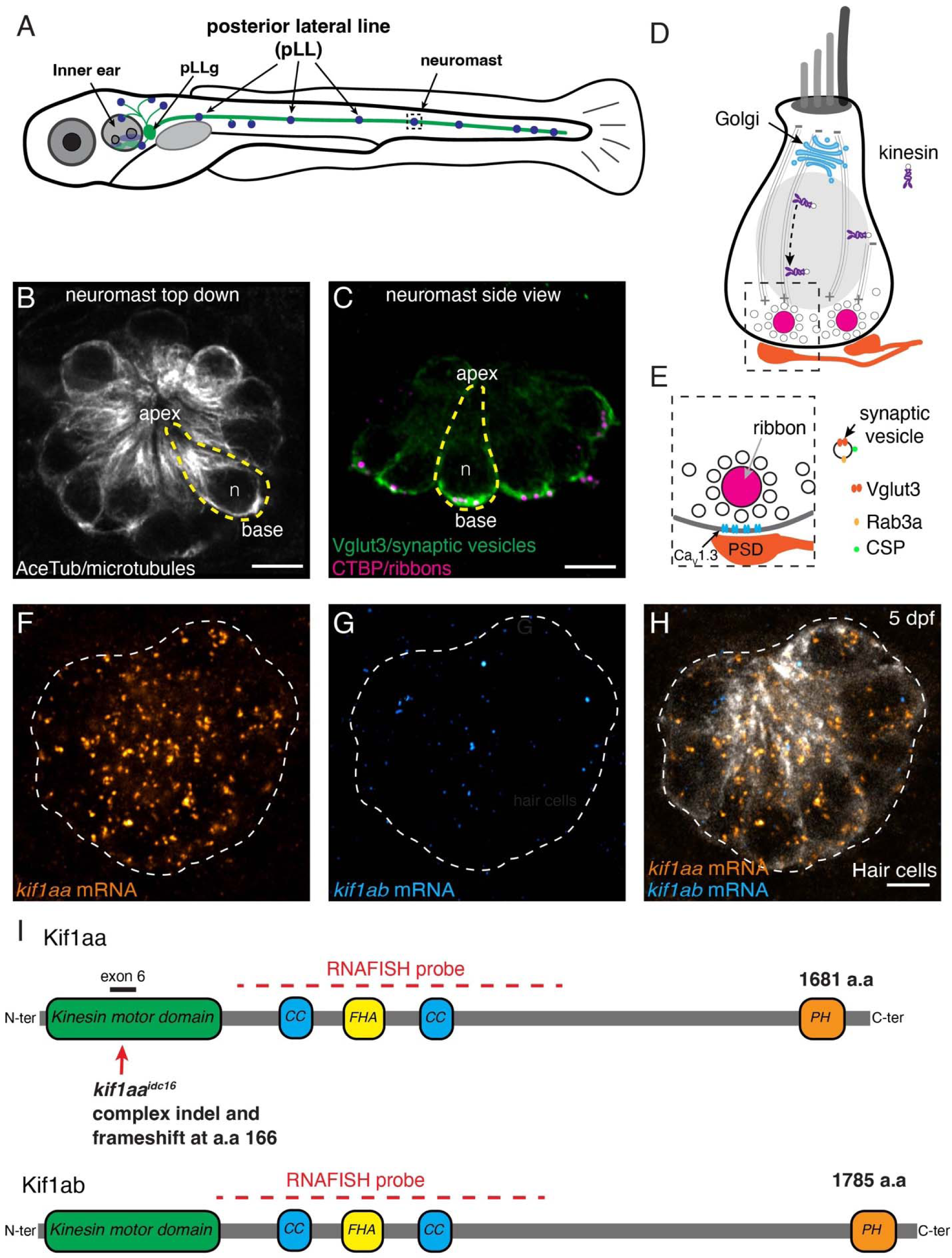
*kif1aa* is expressed in lateral-line hair cells in zebrafish. (**A**) Schematic of a larval zebrafish at 5 dpf. Hair cells are present in the inner ear and lateral line (blue). Hair cells are innervated by neurons in the posterior lateral-line ganglion (pLLg) located near the inner ear (green). (**B-C**) The lateral line is composed of clusters of hair cells called neuromasts (5 dpf). The apex of hair cells project from the center of these clusters, while the ribbon synapses are located at the base of the cells. Neuromasts can be viewed from the top-down **B** or from the side **C**. An individual hair cell in **B** and **C** is outlined in yellow. Within hair cells, microtubule networks extend along the apical-basal axis (**B**). At the base of hair cells, synaptic vesicles (Vglut3 label, green) are enriched near the presynapse or ribbons (CTBP, magenta) (**C**). (**D**) Within hair cells the Golgi is located above the nucleus (gray oval, n in **B**-**C**). The Golgi is where synaptic-vesicle precursors are made *de novo*. Kinesin motors could be used to transport vesicles along microtubules to the cell base. (**E**) Synaptic vesicles surround the presynapse or ribbon in hair cells. Synaptic vesicles contain Rab3a, CSP and Vglut3. Beneath the ribbon Ca_V_1.3 channels are clustered adjacent to the postsynaptic density (PSD). (**F**-**H**) RNA-FISH reveals that *kif1aa* (orange) but not *kif1ab* (cyan) mRNA is present in lateral-line hair cells (YFP, gray) at 5 dpf. (**H**-**I**) Overview of the Kif1aa and Kif1ab proteins and major domains (coiled coil (CC), fork-head associated (FHA), pleckstrin homology (PH)). The location of the germline *kif1aa* lesion in the kinesin motor domain within exon 6 is indicated. The red dashed line indicates the regions encompassed by the RNA-FISH probe. The scale bar in **B**, **C** and **H** = 5 µm.

To study the role of Kif1a and microtubules in synaptic-vesicle transport in hair cells, we examined hair cells in larval zebrafish. In zebrafish, hair cells are found in the inner ear and in the lateral-line system, which are responsible for hearing and balance, and detection of local water flow, respectively (Figure 1A, (Nicolson, 2005; Sheets *et al*., 2021)). In larval zebrafish, the inner ear is composed of 5 sensory organs (3 crista and 2 maculae), while the lateral line is made up of clusters of hair cells called neuromasts that form in lines along the fish. Both of these sensory systems develop rapidly in zebrafish and are mature when larvae are only 5 days post fertilization (dpf) (Zeddies & Fay, 2005; Bhandiwad *et al*., 2013). Importantly, hair cells in the larval zebrafish are morphologically, genetically, and functionally similar to mammalian hair cells, and many of the core synaptic molecules are conserved between zebrafish and mammals. For example, in both mammals and zebrafish: Ribeye makes up the majority of the ribbon density, Vglut3 is the vesicular glutamate transporter required to uptake glutamate into synaptic vesicles, and calcium influx through Ca_v_1.3 channels triggers vesicle fusion (Figure 1E; (Sidi *et al*., 2004; Obholzer *et al*., 2008; Lv *et al*., 2016)). The lateral-line system has proved particularly advantageous to study hair-cell synapses due to the superficial location of neuromasts along the body and the transparency of zebrafish at larval stages. Previous research using both fixed and live imaging approaches in lateral-line organs of larval zebrafish has shown that synaptic vesicles are highly enriched at the base of lateral-line hair cells (Figure 1C, (Obholzer *et al*., 2008; Einhorn *et al*., 2012)). In addition, there are powerful transgenic lines to visualize ribbon synapses or monitor ribbon-synapse activity *in vivo* using genetically-encoded indicators (Zhang *et al*., 2018). To complement this functional imaging, established electrophysiological approaches make it possible to measure vesicle fusion by recording spikes from the afferent neurons that innervate lateral-line neuromasts (Trapani & Nicolson, 2011). Together, these experimental advantages and tools make zebrafish an excellent model to study the role of synaptic-vesicle transport at ribbon synapses in hair cells.

Our study identifies Kif1a as a kinesin motor protein important for transporting and ultimately enriching synaptic vesicles at the presynaptic AZ in hair cells. We show that while both zebrafish ohnologues (paralogous sequences that result from complete genome duplication) of mammalian *Kif1a*, *kif1aa* and *kif1ab,* are expressed in inner-ear hair cells, only *kif1aa* is expressed in lateral-line hair cells. We find that *kif1aa* mutants are unable to enrich synaptic vesicles at the presynaptic AZ of lateral-line hair cells. Likewise, wild-type zebrafish exhibit the same loss of synaptic-vesicle enrichment when microtubules are destabilized. Functionally, we show that *kif1aa* mutants have normal evoked presynaptic-calcium responses, but that postsynaptic-calcium responses in afferent neurons are dramatically reduced. Further, the spontaneous spike rate of lateral-line afferent neurons is dramatically decreased in *kif1aa* mutants. Behaviorally, we find that *kif1aa* mutants have impaired station holding behavior during rheotaxis–a behavior that depends on a functional lateral-line system. Overall, our work demonstrates that a Kif1a-based mechanism is essential to enrich synaptic vesicles at ribbon synapses in hair cells. This enrichment is essential for ribbon-synapse function and ultimately for sensory behavior.

## Results

### Kif1aa is expressed in lateral-line hair cells and impacts ribbon-synapse counts

Work in neurons has demonstrated that Kif1a is an anterograde kinesin motor that transports synaptic-vesicle precursors towards the plus ends of microtubules (Okada *et al*., 1995). Recent work in both mouse and zebrafish hair cells has demonstrated that microtubules are polarized, with their plus ends growing toward the cell base, near the presynaptic AZ (Figure 1B; (Hussain *et al*., 2024; Voorn *et al*., 2024)). Further, RNAseq data indicates robust levels of *Kif1a* transcripts in mouse auditory hair cells and zebrafish hair cells (Lush *et al*., 2019; Kolla *et al*., 2020; Sur *et al*., 2023). Based on work in neurons, we used zebrafish to test whether Kif1a is a kinesin motor involved in the transport of synaptic vesicles to ribbon synapses in hair cells.

In zebrafish, there are two ohnologues of mammalian *Kif1a*, *kif1aa* and *kif1ab* (Figure 1I). Kif1aa and Kif1ab proteins show a high degree of identity to each other (89 %, NCBI BLAST) and to human KIF1A (84 and 85 % identity respectively, NCBI BLAST). Recent scRNAseq work has detected widespread expression of both *kif1aa* and *kif1ab* transcripts throughout the zebrafish nervous system (Sur *et al*., 2023). Similarly, widespread expression of *Kif1a* in the nervous system of mice is also well-documented (Okada *et al*., 1995). We used high-resolution RNA fluorescence in situ hybridization (RNA-FISH) to verify whether the *kif1aa* and *kif1ab* mRNAs are present in zebrafish hair cells (Choi et al., 2018). We found that only *kif1aa* mRNA was present in hair cells of the zebrafish lateral line (Figure 1F-H). In contrast, both *kif1aa* and *kif1ab* mRNA were present in the hair cells of the zebrafish inner ear (Figure 1-S1). In the inner ear we observed both *kif1aa* and *kif1ab* mRNA in the cristae (which detects angular acceleration), the anterior macula (which detects gravity and is required for balance), and the posterior macula (which primarily detects auditory stimuli) (Fay & Popper, 2000).

After validating that *Kif1a* ohnologues are indeed expressed in hair cells, we leveraged zebrafish genetics to test whether Kif1a plays a role in the transport of synaptic vesicles in hair cells. We focused our efforts on a germline *kif1aa* mutant we created in a previous study (Hussain *et al*., 2024). This mutant has a stop codon in the Kif1aa motor domain and is predicted to be a null mutation (Figure 1I). This mutant should impair Kif1a in lateral-line hair cells but leave Kif1a function at least partially intact in the inner ear and the nervous system, due to the expression of the ohnologue *kif1ab*. We found via RNA-FISH that *kif1aa* mRNA expression was reduced by ∼60 % in lateral-line hair cells in *kif1aa* mutants compared to sibling controls (Figure 1-S2, control: 403.3 ± 65.8 puncta; *kif1aa*: 172.7 ± 48.6 puncta, n = 18 control and 16 *kif1aa* neuromasts, unpaired t-test, p < 0.0001). Overall, this reduction in mRNA levels provides evidence that we have disrupted the *kif1aa* locus and that there is significant nonsense-mediated degradation of *kif1aa* mRNA in the lateral-line hair cells of *kif1aa* mutants.

Before assessing synaptic vesicles in *kif1aa* mutant hair cells, we first performed a gross assessment of lateral-line neuromasts using immunohistochemistry to label hair cells and ribbon synapses (Figure 2A-B). We performed our analyses at 5 dpf, when the majority of the hair cells are mature, and the lateral-line system is functional. We used antibodies against Myosin7a (Myo7a) to label hair cells, along with Ribeye b (Rib b) and Membrane-associated guanylate kinase (MAGUK) to label the pre- and postsynapses respectively. We found that *kif1aa* mutants have a similar number of hair cells per neuromast compared to sibling controls (Figure 2C, control: 15.6 ± 1.3; *kif1aa*: 16.0 ± 1.7, n = 18 control and 15 *kif1aa* neuromasts; unpaired t-test, p = 0.399). For our synapse analysis we examined 2D maximum intensity projections and quantified the number and area of ribbons and postsynapses. Similar to our previous work on developing hair cells (3 dpf), we found that there were significantly fewer complete synapses (paired Rib b-MAGUK puncta) per hair cell in mature lateral-line hair cells (5 dpf) in *kif1aa* mutants compared to sibling controls (Figure 2D, control: 3.13 ± 0.35; *kif1aa*: 2.68 ± 0.14, n = 18 control and *kif1aa* 15 neuromasts; unpaired t-test, p = 0.00870). Quantification of pre- and post-synapse size at complete synapses revealed no difference in presynapse size, but slightly larger postsynapses in *kif1aa* mutants compared to controls (Figure 2E, presynapse area, control: 0.21 ± 0.02; *kif1aa*: 0.20 ± 0.02, unpaired t-test, p = 0.345; Figure 2F, postsynapse area, control: 0.19 ± 0.02; *kif1aa*: 0.20 ± 0.02, unpaired t-test, p = 0.0498). Overall, we found that *Kif1a* ohnologues are present in hair cells in zebrafish. In the lateral-line, loss of Kif1aa does not alter hair-cell counts but does result in slightly fewer ribbon synapses.

**Figure 2.**
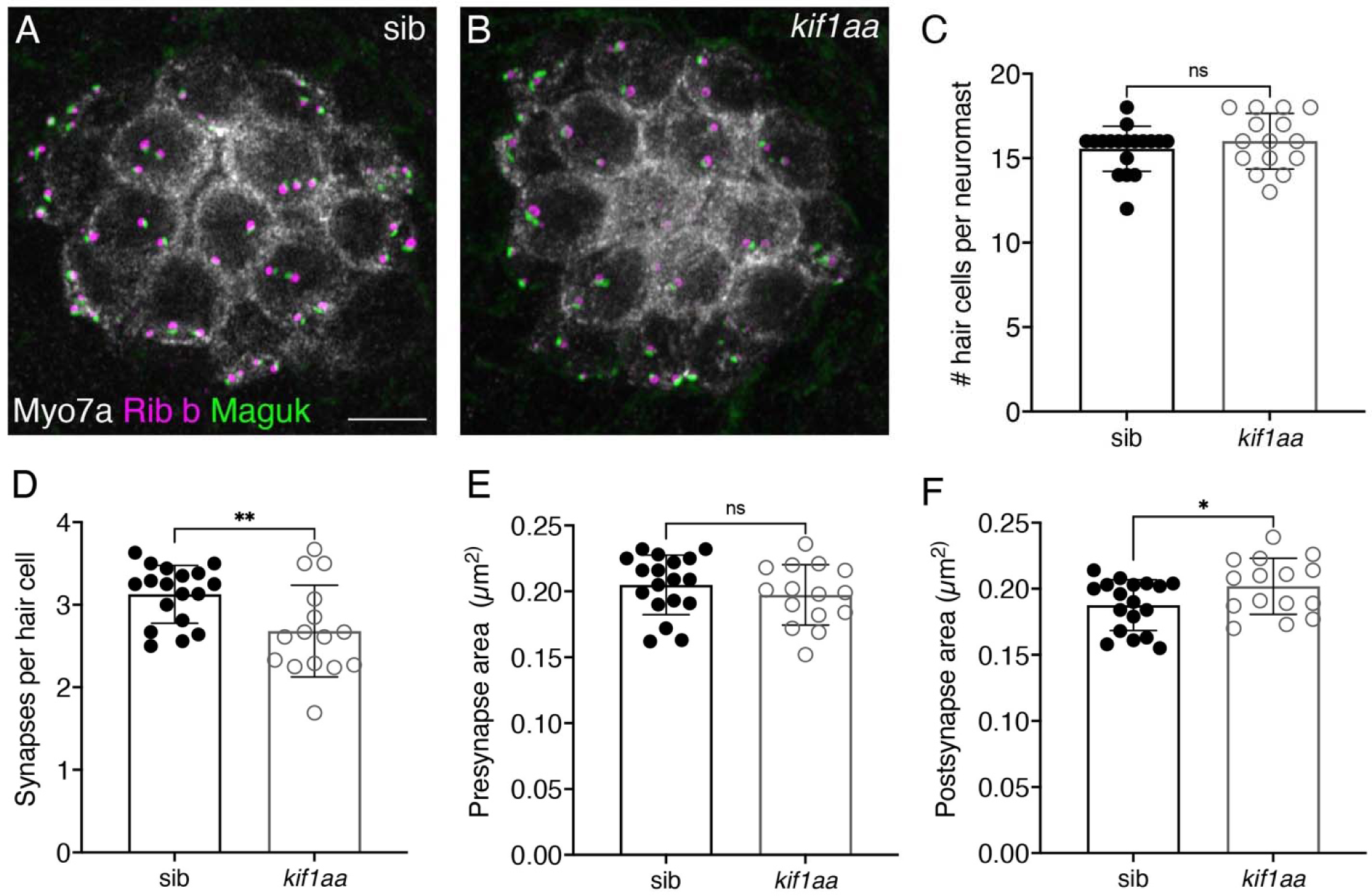
*Kif1aa* mutants have fewer ribbon synapses. (**A-B**) Example immunolabel of neuromasts at 5 dpf in a *kif1aa* mutant (**B**) or sibling control (**A**). Myosin7a (Myo7a) labels hair cells, Ribeye b (Rib b) labels ribbons or presynapses, and Maguk labels postsynapses. (**C**) The number of hair cells per neuromast is the same in *kif1aa* mutants compared to sibling controls (control: 15.6 ± 1.3; *kif1aa*: 16.0 ± 1.7, unpaired t-test, p = 0.399). (**D**) *Kif1aa* mutants have fewer complete synapses per cell (control: 3.13 ± 0.35; *kif1aa*: 2.68 ± 0.56, unpaired t-test, p = 0.00870). (**E**) In *kif1aa* mutants the average area of Rib b puncta (presynaptic) was similar to sibling controls (control: 0.21 µm^2^ ± 0.023; *kif1aa:* 0.20 µm^2^ ± 0.02, unpaired t-test, p = 0.345). (**F**) In *kif1aa* mutants the average area of Maguk puncta (postsynaptic) was slightly larger compared to sibling controls (control: 0.19 µm^2^ ± 0.02; *kif1aa:* 0.20 µm^2^ ± 0.02, unpaired t-test, p = 0.0498). n = 18 control and 15 *kif1aa* neuromasts in **C-F**. Scale bar in **A** = 5 µm.

### Kif1aa is essential for synaptic-vesicle distribution in hair cells

Our overall assessment suggests that in *kif1a* mutants, ribbon synapses are largely intact, although fewer in number. As previous work in neurons has shown that Kif1a transports synaptic-vesicle precursors to the presynaptic AZ, we assessed whether loss of Kif1aa impacts synaptic-vesicle distribution at the presynapse in hair cells. We imaged both live and fixed preparations to visualize synaptic-vesicle distributions in hair cells of *kif1aa* mutants. To label synaptic vesicles live, we used the vital dye LysoTracker. This dye has been shown to label acidified organelles, including synaptic vesicles in lateral-line hair cells (Einhorn *et al*., 2012). To label synaptic vesicles in fixed samples, we used immunohistochemistry to label Vglut3, the vesicular glutamate transporter that labels synaptic vesicles in hair cells of zebrafish and mice (Obholzer *et al*., 2008; Schraven *et al*., 2012)

In line with previous work, we observed an enrichment of LysoTracker dye at the cell base near the presynaptic AZ in sibling controls (Figure 3A,C; (Einhorn *et al*., 2012)). In contrast, we did not observe enrichment of LysoTracker at the base of hair cells in *kif1aa* mutants (Figure 3B,C). To quantify Lysotracker enrichment, we measured the base to apex ratio of fluorescence in individual hair cells (Figure 3C). Using this metric, we found that *kif1aa* mutants enrich significantly less LysoTracker at the cell base (Figure 3D, control: 7.38 ± 1.16; *kif1aa*: 0.52 ± 0.16 n = 10 control and *kif1aa* neuromasts, unpaired t-test, p<0.0001). Interestingly, we observed that the base to apex ratio was less than 1 in Kif1aa-deficient hair cells. This indicates that the LysoTracker label is more enriched in the hair-cell apex in *kif1aa* mutants.

**Figure 3.**
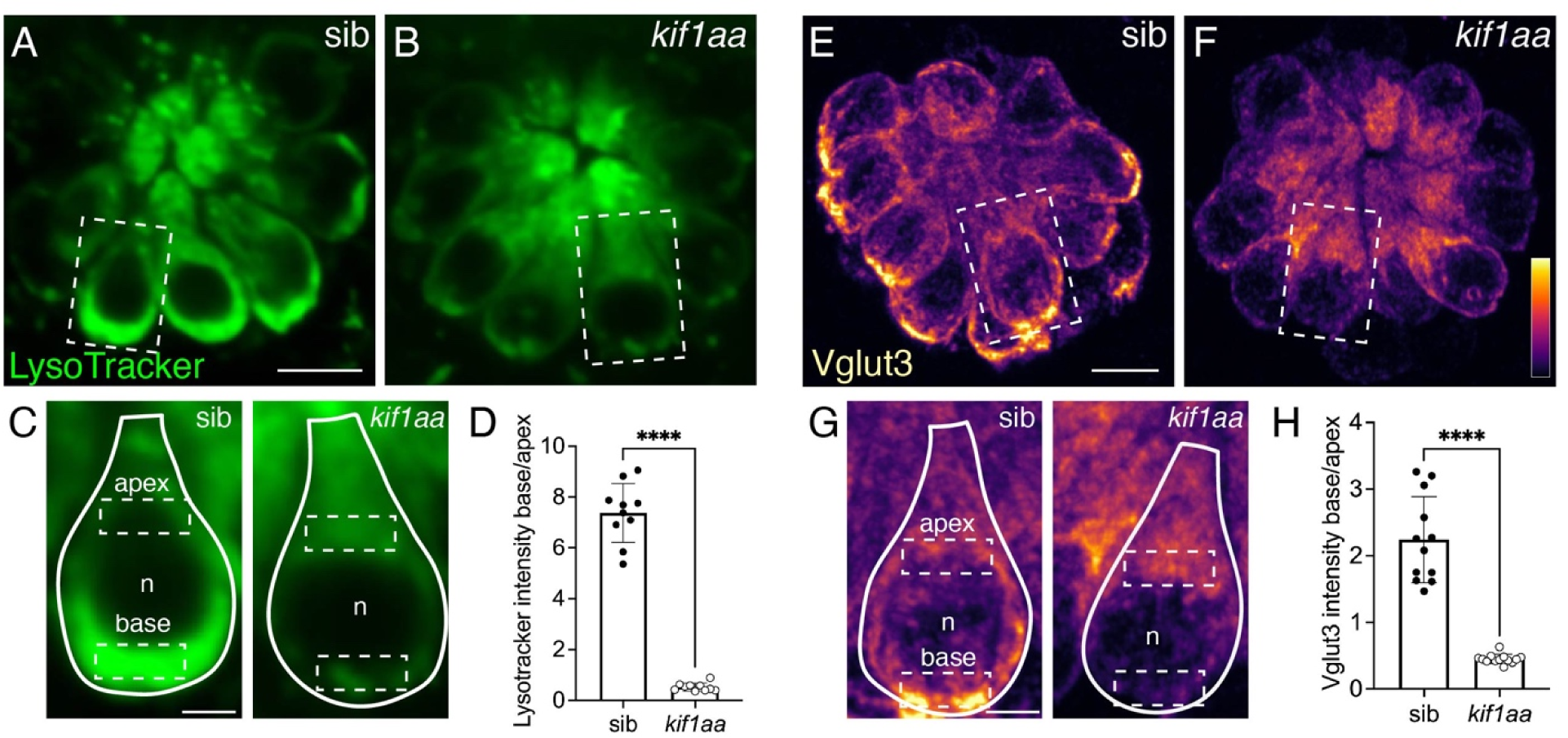
*Kif1aa* mutants enrich less LysoTracker and Vglut3 at the presynapse. (**A-C**) Example live image of Lysotracker Red (green) to label synaptic vesicles in neuromasts at 5 dpf in a *kif1aa* mutant (**B**) or sibling control (**A**). The dashed box in each images indicates the hair cells magnified and outlined with a solid line in (**C**). n indicates nucleus, and dashed boxes indicate example ROIs of the apical and basal regions used for intensity analysis in **D**. (**D**) Quantification of LysoTracker shows that in *kif1aa* mutants there is significantly less enrichment at the cell base compared to sibling controls (control: 7.38 ± 1.16; *kif1aa*: 0.52 ± 0.16, n = 10 *kif1aa* and control neuromasts, unpaired t-test, p<0.0001). (**E-G**) Example immunostain of Vglut3 to label synaptic vesicles in neuromasts at 5 dpf in a *kif1aa* mutant (**F**) or sibling control (**E**). The dashed box in each images indicates the hair cells magnified and outlined with a solid line in (**G**). n indicates nucleus, and dashed boxes indicate example ROIs of the apical and basal regions used for intensity analysis in **H**. (**H**) Quantification reveals that Vglut3 is significantly less enriched at the cell base in *kif1aa* mutants compared to sibling controls (control: 2.24 ± 0.65; *kif1aa*: 0.45 ± 0.07, n = 12 control and 13 *kif1aa* neuromasts, unpaired t-test, p<0.0001) Scale bar in **A** and **E** = 5 µm and 2 µm in **C** and **G**.

Similar to LysoTracker label, and consistent with previous work in lateral-line hair cells, we observed that Vglut3 immunolabel was enriched at the cell base in sibling controls (Figure 3E,G; (Obholzer *et al*., 2008)). This enrichment was not observed in *kif1aa* mutants (Figure 3F,G). We quantified enrichment by measuring the base to apex ratio of Vglut3 immunolabel. We found that the base to apex ratio of Vglut3 label was significantly reduced in *kif1aa* mutants compared to sibling controls (Figure 3H, control: 2.24 ± 0.65; *kif1aa*: 0.45 ± 0.07; n = 12 control and 13 *kif1aa* neuromasts, unpaired t-test, p<0.0001).

To further verify that *kif1aa* mutants fail to enrich synaptic vesicles at the base of lateral-line hair cells, we used immunohistochemistry to label Rab3a and Cysteine string protein (CSP) (Figure 3-S1A-C and E-G). Rab3a and CSP are synaptic-vesicle markers that have also been shown to be enriched at the base of lateral-line hair cells (Einhorn *et al*., 2012; Kindt & Sheets, 2018). We measured the base to apex ratio of Rab3a and CSP immunolabels to quantify synaptic-vesicle enrichment. We observed that similar to our Vglut3 immunolabel, the base to apex ratio of both Rab3a and CSP immunolabels were significantly reduced in *kif1aa* mutants compared to sibling controls (Figure 3-S1D, CSP, control: 1.86 ± 0.34; *kif1aa*: 0.58 ± 0.07; n = 10 control and *kif1aa* neuromasts, unpaired t-test, p<0.0001 and Figure 3-S1H, Rab3a, control: 1.57 ± 0.30; *kif1aa*: 0.63 ± 0.15; n = 9 control and *kif1aa* neuromasts, unpaired t-test, p<0.0001).

In addition to lateral-line hair cells, we also examined synaptic-vesicle distribution in inner-ear hair cells in *kif1aa* mutants. For our analysis, we focused on Vglut3 labeling, as Vglut3 has been shown to be essential for both inner ear and lateral-line function in zebrafish (Obholzer *et al*., 2008). First, we examined Vglut3 distribution in cristae. Interestingly, in sibling controls, we observed that Vglut3 label was only enriched in a subset of the hair cells (Figure 3-S2A, blue outlines). Previous work has shown that zebrafish cristae have two main hair-cell types that can be distinguished morphologically as “tall” and “short” hair cells (Zhu *et al*., 2020; Smith *et al*., 2020). In sibling controls, we found that the tall cells enriched Vglut3 at the cell base. Compared to tall cells, short cells had nearly undetectable levels of Vglut3 (Figure 3-S2A, yellow outlines). A difference in Vglut3 immunolabel has also been observed between hair-cell types in mouse vestibular hair cells where type 2 but not type 1 vestibular hair cells express Vglut3 (Schraven *et al*., 2012). Importantly, when we examined *kif1aa* mutants we found that tall cells fail to enrich Vglut3 immunolabel at the cell base (Figure 3-S2B, blue outlines). This observation in *kif1aa* mutants indicates that *kif1ab* mRNA in cristae is not sufficient for normal Vglut3 enrichment in *kif1aa* mutants (Figure 1-S1B). In addition to cristae, we also examined Vglut3 immunolabel in the anterior and posterior macula. Unlike in the cristae, we found that in sibling controls all macular hair cells had detectable amounts of Vglut3 immunolabel and showed enrichment of Vglut3 at the cell base (Figure 3-S2C). In general, Vglut3 immunolabel was qualitatively lower yet still enriched at the cell base in the macular hair cells of *kif1aa* mutants (Figure 3-S2D). This indicates that Kif1aa is not essential for normal synaptic-vesicle distribution in macular hair cells.

Overall, our live and fixed imaging of synaptic-vesicle distribution in *kif1aa* mutants indicates that Kif1aa is essential to enriching synaptic vesicles at the presynaptic AZ of lateral-line hair cells and specifically in tall cells of the cristae. In contrast, loss of Kif1aa has a more moderate impact on synaptic-vesicle distribution in macular hair cells.

### An intact microtubule network is essential for synaptic-vesicle distribution

Kif1a is known to transport synaptic-vesicle precursors towards the plus ends of microtubules. Therefore, we employed a pharmacological approach to understand the role that an intact microtubule network plays in synaptic-vesicle distribution in lateral-line hair cells. Nocodazole has been shown to block the self-assembly of tubulin and depolymerize preassembled microtubules (Samson *et al*., 1979). In addition, recent work has shown that treatment with 250-500 nM nocodazole can disrupt microtubule networks in developing lateral-line hair cells (Hussain *et al*., 2024). We found that incubating 5 dpf larvae in 250 nM nocodazole for 2 hrs disrupted microtubules in mature hair cells without any detectable hair-cell death (Figure 4-S1). We used nocodazole along with LysoTracker label or Vglut3 immunolabel to assess the impact of microtubule disruption on synaptic-vesicle distribution.

**Figure 4.**
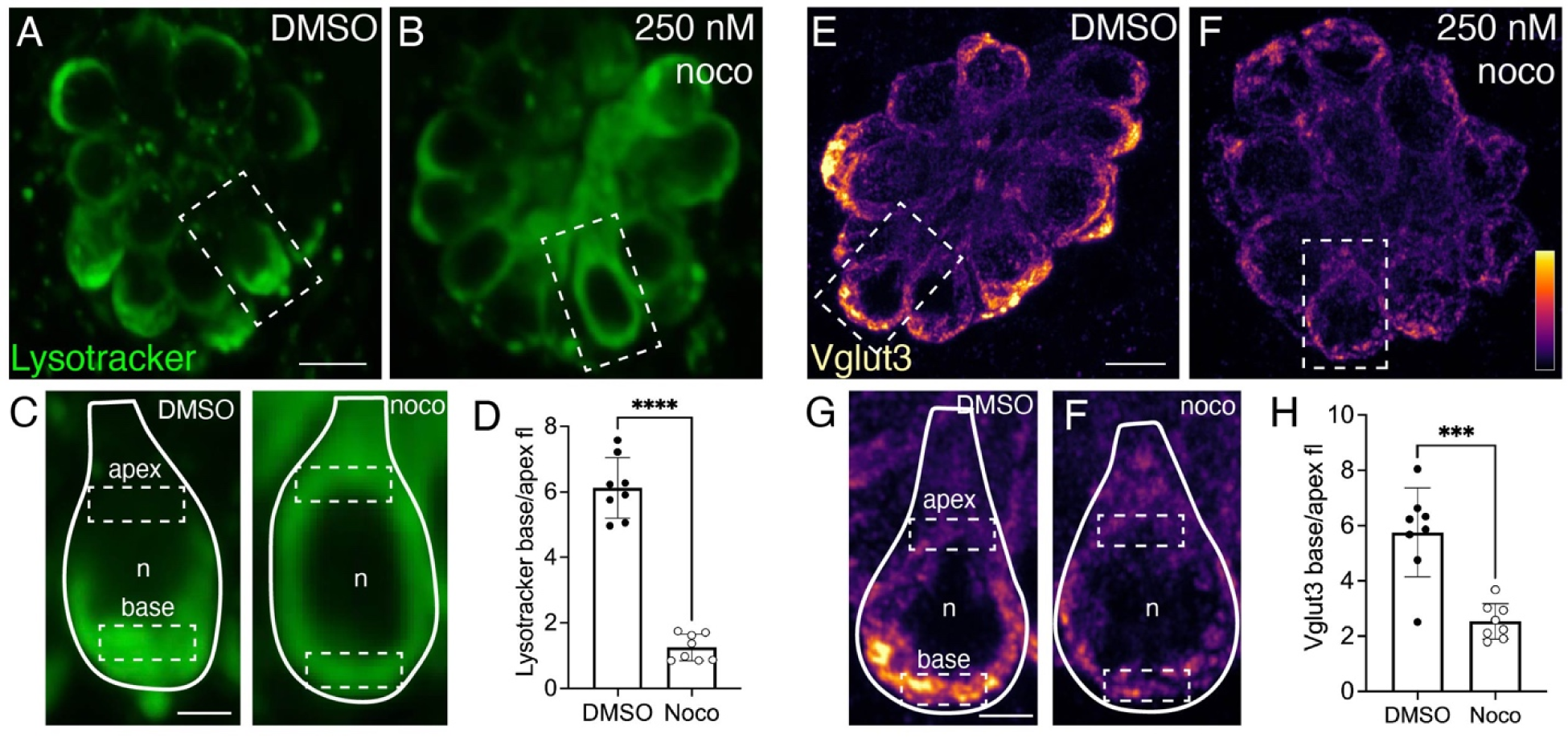
An intact microtubule network is required to enrich LysoTracker and Vglut3 at the presynapse. (**A-C**) Example live image of LysoTracker Red (green) to label synaptic vesicles in neuromasts at 5 dpf in wild-type larva treated with 250 nM nocodazole (**B**) or DMSO control (**A**). The dashed box in each image indicates the hair cell magnified and outlined with a solid line in (**C**). n indicates nucleus, and dashed boxes indicate example ROIs of apical and basal regions used for the analysis in **D**. (**D**) Quantification shows significantly less LysoTracker enrichment at the cell base in nocodazole-treated larvae compared to DMSO control (control: 6.12 ± 0.93; nocodazole: 1.25 ± 0.40, n = 8 control and nocodazole neuromasts, unpaired t-test, p<0.0001). (**E-G**) Example immunolabel of Vglut3 to label synaptic vesicles in neuromasts at 5 dpf in larva treated with 250 nM nocodazole (**F**) or DMSO control (**E**). The dashed box in each image indicates the hair cell magnified and outlined with a solid line in (**G**). n indicates nucleus, and dashed boxes indicate example ROIs of the apical and basal regions used for intensity analysis in **H**. (**H**) Quantification reveals significantly less Vglut3 enrichment at the cell base in nocodazole-treated larvae compared to DMSO controls (control: 5.75 ± 1.61; nocodazole: 2.53 ± 0.65, n = 8 control and nocodazole neuromasts, unpaired t-test, p<0.0001). Scale bar in **A** and **E** = 5 µm and 2 µm in **C** and **G**.

After incubation with 250 nM nocodazole, we observed less basal enrichment of Lysotracker compared to DMSO controls (Figure 4A-C). To quantify label enrichment, we measured the base to apex ratio of Lysotracker fluorescence in hair cells (Figure 4C-D). We found significantly less basal enrichment of Lysotracker in nocodazole-treated hair cells compared to DMSO controls (Figure 4D, control: 6.12 ± 0.93; nocodazole: 1.25 ± 0.40 n = 8 control and nocodazole neuromasts, unpaired t-test, p<0.0001). In addition, we found significantly less basal enrichment of Vglut3 in hair cells in nocodazole-treated larvae compared to DMSO controls (Figure 4H-E, control: 5.75 ± 1.61; nocodazole: 2.53 ± 0.65; n = 8 control and nocodazole neuromasts, unpaired t-test, p<0.0001). Together, our nocodazole treatments indicate that an intact microtubule network and functional Kif1aa are necessary to localize synaptic vesicles at the presynapse in lateral-line hair cells.

### Kif1aa is important to maintain synaptic vesicles at ribbon synapses

Our live and fixed confocal imaging strongly suggests that synaptic vesicles fail to enrich at the cell base in *kif1aa* mutants (Figure 3, Figure 3-S1, S2). To understand how synaptic vesicles are impacted specifically at sites of release we examined synaptic vesicles at individual ribbons in lateral-line hair cells. We first examined our LysoTracker label more closely at individual ribbons. To detect ribbons within the LysoTracker label, we used a transgenic line that labels ribbons *Tg(myo6b:Rib a-TagRFP).* In addition to LysoTracker measurements, we also used transmission electron microscopy (TEM) to visualize individual synaptic vesicles at ribbons.

In control hair cells, we observed an intense LysoTracker label or halo surrounding individual ribbons (Figure 5A-B). This halo likely represents the population of synaptic vesicles tethered to the ribbon and is comparable to previous observations in lateral-line hair cells (Einhorn et al, 2012). We quantified the amount of LysoTracker in ROIs directly on individual ribbons. We measured Lysotracker intensity in a single place that encompassed the center of the ribbon. Using this approach, we found that the amount of Lysotracker at individual ribbons was significantly reduced in *kif1aa* mutants compared to sibling controls (Figure 5C, control: 2582 ± 450; *kif1aa*: 1810 ± 650, n = 18 control and *kif1aa* neuromasts, unpaired t-test, p = 0.0003). This indicates that there is less Lysotracker label and likely fewer synaptic vesicles at ribbons in *kif1aa* mutants.

**Figure 5.**
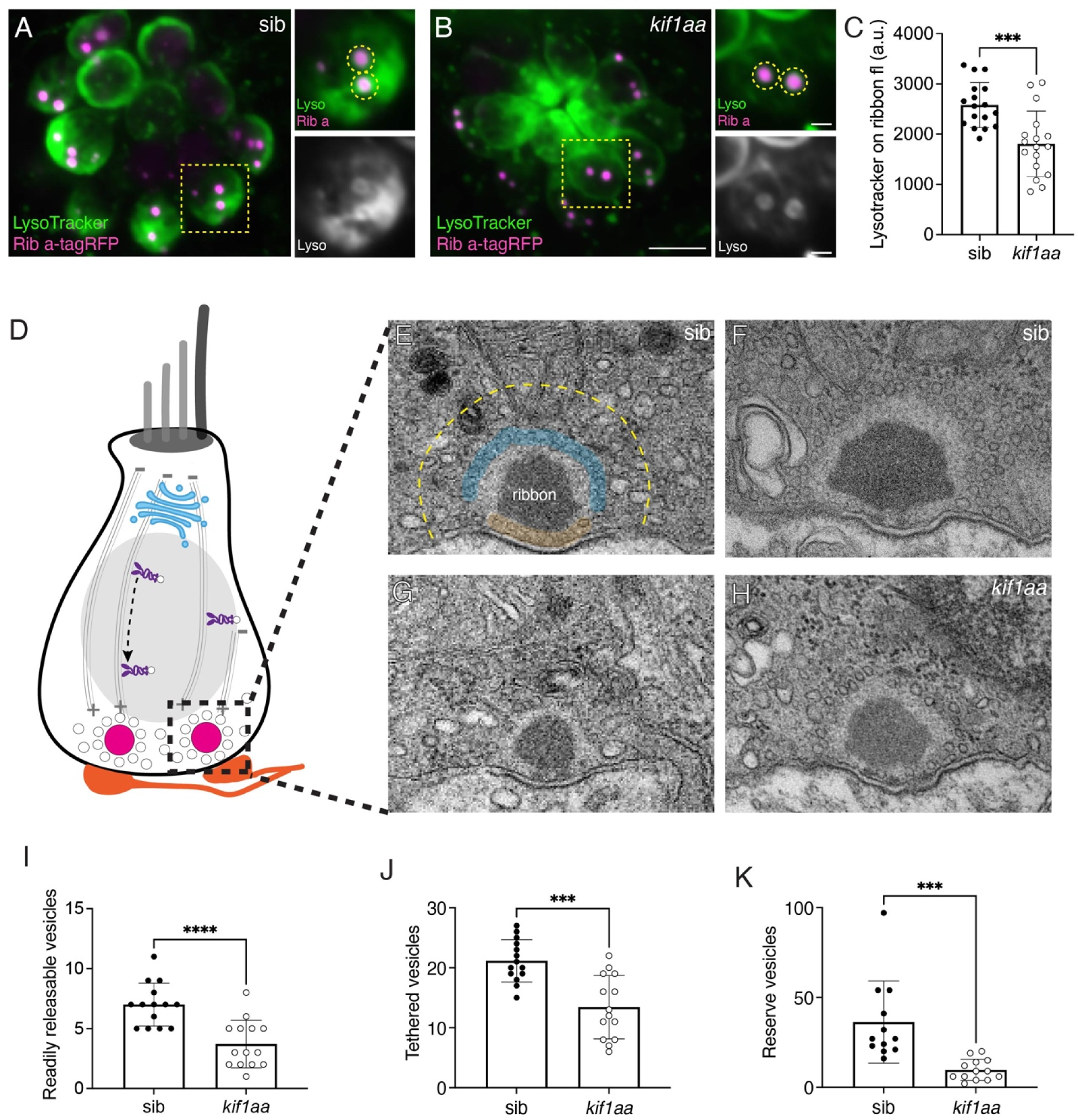
Kif1a is important to maintain synaptic vesicles at ribbon synapses. (**A-B**) Example live images of neuromasts at 5 dpf in a *kif1aa* mutant (**B**) or sibling control (**A**) at 5 dpf. Synaptic vesicles are labeled with LysoTracker Green (green), and ribbons are labeled with Rib a-TagRFP (magenta). The yellow dashed lines in **A** and **B** were used to create the insets shown to the side. The top inset image shows the base of this single hair cell labeled with Lysotracker (green) and Rib a-TagRFP (magenta), while the bottom inset shows just LysoTracker (gray). Insets are only a partial projection (single cell) of the larger image. The yellow dashed circles indicated ROIs used to quantify LysoTracker intensity at ribbons. (**C**) The average LysoTracker intensity at ribbons is significantly lower in *kif1aa* mutants compared to sibling controls (control: 2582 ± 450; *kif1aa*: 1810 ± 650, n = 17 control and *kif1aa* neuromasts, unpaired t-test, p = 0.0003). (**D**) Schematic of an individual hair cell with a ribbon (black dashed box) like ones examined for TEM. (**E-H**) Representative TEM images of ribbon synapses from *kif1aa* mutants (**G-H**) or wild-type sibling controls (**E-F**) at 5 dpf. In **E “**tethered vesicles” are in blue, “readily releasable vesicles” in orange, while “the reserve pool” within 200 nm of the ribbon is encompassed by the yellow dashed line. (**I-K**) All synaptic vesicle pools are reduced in *kif1aa* mutants compared to sibling controls (**I**, readily releasable, control: 7 ± 1.8; *kif1aa*: 3.7 ± 2.0, unpaired t-test, p<0.0001; **J**, tethered, control: 21.1 ± 3.5; *kif1aa*: 13.4 ± 5.3, unpaired t-test, p = 0.0001; **K**, reserve pool, control: 36.3 ± 23.0; *kif1aa*: 9.7 ± 5.9, unpaired t-test, p = 0.0005). n = 14 control *kif1aa* ribbons in **I-J,** and n = 12 control and 13 *kif1aa* ribbons in **K**. Scale bar in **A** = 5 µm, inset is 0.1 µm, **F** = 250 nm.

To quantify synaptic vesicles more definitively, we used transmission electron microscopy (TEM) to visualize ribbons in lateral-line hair cells. We only examined micrographs that contained images of a centrally sectioned ribbon adjacent to a well-defined postsynaptic density (Figure 5D-H). We quantified three main populations of synaptic vesicles localized in the vicinity of ribbons. We define these populations as following: 1) “vesicles tethered to the ribbon”; 2) “ready releasable vesicles” located directly beneath the ribbon; and 3) “reserve pool of vesicles” residing within 200 nm of the ribbon (Figure 5E, zones shaded in blue and orange, and a yellow dashed line) (Lenzi *et al*., 1999; Nouvian *et al*., 2006). We found that all three of these synaptic-vesicle populations were reduced in *kif1aa* mutants compared to sibling controls (Figure 5I, readily releasable, control: 7 ± 1.8; *kif1aa*: 3.7 ± 2.0, unpaired t-test, p<0.0001; Figure 5J, tethered, control: 21.1 ± 3.5; *kif1aa*: 13.4 ± 5.3, unpaired t-test, p = 0.0001; Figure 5K, reserve pool, control: 36.3 ± 23.0; *kif1aa*: 9.7 ± 5.9, unpaired t-test, p = 0.0005). Our TEM quantification of synaptic vesicles is consistent with our LysoTracker labeling at ribbons, indicating that LysoTracker can be a quantifiable way to measure synaptic vesicles at ribbons using confocal microscopy. Overall, using both of these methods, we find that synaptic vesicles are depleted at ribbons in *kif1aa* mutants.

### Loss of Kif1aa does not impact mechanosensitive or presynaptic calcium responses

Our results indicate that loss of Kif1aa in lateral-line hair cells dramatically decrease synaptic vesicle localization at the hair-cell base and at ribbon synapses (Figures 3, 5 and Figure 3-S1). But what impact this impairment in localization has on synaptic-vesicle release at ribbon synapses was unclear. Before examining vesicle release, performed several control experiments in *kif1aa* mutants and sibling controls.

Prior to assessing vesicle release in lateral-line hair cells, we first examined the localization of voltage-dependent calcium channels (Ca_V_1.3). Ca_V_1.3 channel localization can impact presynaptic calcium responses as well as both spontaneous and evoked vesicle fusion (Brandt *et al*., 2005; Trapani & Nicolson, 2011). In addition, calcium channels have been shown to be transported in synaptic precursor vesicles (Petzoldt, 2023). We used immunohistochemistry to examine Ca_V_1.3 clusters and Rib b to label ribbons (Figure 6A-B). For our analyses we examined maximum intensity projections and quantified the number of puncta, along with the 2D area, average intensity, and integrated density of Ca_V_1.3 puncta. We first quantified the number of paired Ca_V_1.3-Rib b puncta and found that there were significantly fewer paired puncta per hair cell in *kif1aa* mutants compared to sibling controls (Figure 6C, control: 4.13 ± 0.39; *kif1aa*: 3.59 ± 0.41; n = 13 control and 9 *kif1aa* neuromasts, unpaired t-test, p = 0.00550). Fewer paired Ca_V_1.3-Rib b puncta per cell is consistent with an overall reduction in the number of complete synapses per cell we observed in *kif1aa* mutants (Figure 2D). Looking more closely at Ca_V_1.3 puncta, we did not observe a significant difference in the average size of Ca_V_1.3 puncta between sibling controls and *kif1aa* mutants (Figure 6D, control: 0.14 µm^2^ ± 0.02; *kif1aa*: 0.15 µm^2^ ± 0.02; n = 13 control and 9 *kif1aa* neuromasts, unpaired t-test, p = 0.0953). However, both the mean intensity and integrated intensity of Ca_V_1.3 puncta were significantly higher in *kif1aa* mutants compared to controls (Figure 6E, mean intensity, control: 1685 ± 459; *kif1aa*: 2961 ± 1002, unpaired t-test, p = 0.000600; Figure 6F, integrated intensity, control: 229.0 ± 73.6; *kif1aa*: 444.1 ± 180.2, unpaired t-test, p = 0.000900; n = 13 control and 9 *kif1aa* neuromasts). Together, our Ca_V_1.3 immunolabeling experiments indicate that although there are fewer paired Ca_V_1.3-Rib b puncta in *kif1aa* mutants, on average more Ca_V_1.3 channels may reside within each Ca_V_1.3 puncta.

**Figure 6.**
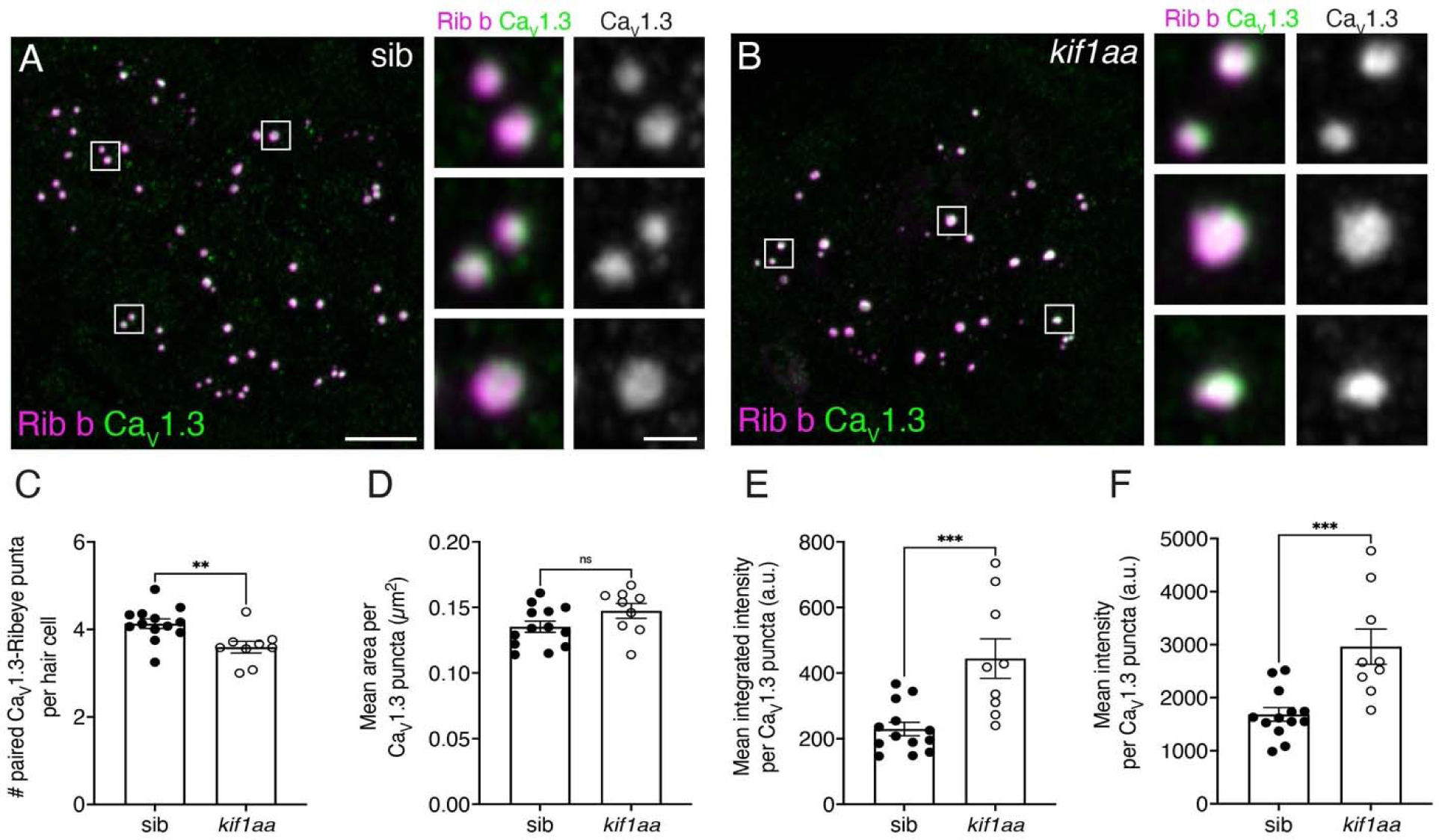
There are fewer paired Cav1.3-Rib b puncta in *kif1aa* mutants. (**A-B**) Example immunostain of neuromasts at 5 dpf in *kif1aa* mutants (**B**) or sibling control (**A**). Ribeye b (Rib b) labels ribbons or presynapses, and Ca_V_1.3 labels calcium channels. Insets to the side show high magnification images of individual synapses. (**C**) The number of paired Rib b-Ca_V_1.3 puncta is significantly reduced in *kif1aa* mutants compared to sibling controls (control: 4.13 ± 0.39; *kif1aa*: 3.59 ± 0.41; n = 13 control and 9 *kif1aa* neuromasts, unpaired t-test, p = 0.00550). (**D**) The average size of Ca_V_1.3 puncta was similar between sibling controls and *kif1aa* mutants (control: 0.14 ± 0.02; *kif1aa*: 0.15 ± 0.02, unpaired t-test, p = 0.0953). (**E-F**) The mean intensity and integrated intensity of Ca_V_1.3 puncta were significantly higher in *kif1aa* mutants compared to controls (**E**, mean intensity, control: 1685 ± 459; *kif1aa*: 2961 ± 1002, unpaired t-test, p = 0.000600; **F**, integrated intensity, control: 229.0 ± 73.6; *kif1aa*: 444.1 ± 180.2, unpaired t-test, p = 0.0009). n = 13 control and 9 *kif1aa* neuromasts in **C-F**. Scale bar in **A** = 5 µm, inset is 0.5 µm.

After examining Ca_V_1.3 channel localization at ribbon synapses we also did control experiments to assess hair-cell function. We first examined mechanosensation and presynaptic calcium responses. These are two activity-dependent events in hair cells that are required to drive synaptic vesicle fusion at ribbon synapses. To assess hair-cell mechanosensation we used two independent methods: FM 1-43 labeling and calcium imaging. Numerous studies have demonstrated that the vital dye FM 1-43 is a straightforward way to assess whether hair cells have intact mechanotransduction channels (Gale *et al*., 2001; Meyers *et al*., 2003). When applied to the media zebrafish are housed in, FM 1-43 enters mechanotransduction channels in lateral-line hair cells in seconds (Seiler & Nicolson, 1999). In contrast, mutants lacking mechanotransduction fail to label with FM 1-43. When we applied FM 1-43 to *kif1aa* mutants we found similar intensities of label compared to sibling controls (Figure 7-S1A-C, control: 2182 ± 227.1; *kif1aa*: 2034 ± 302.8, n = 8 control and 9 *kif1aa* neuromasts, unpaired t-test, p = 0.276). Normal FM 1-43 label in *kif1aa* mutants indicates that mechanosensation is largely intact.

**Figure 7.**
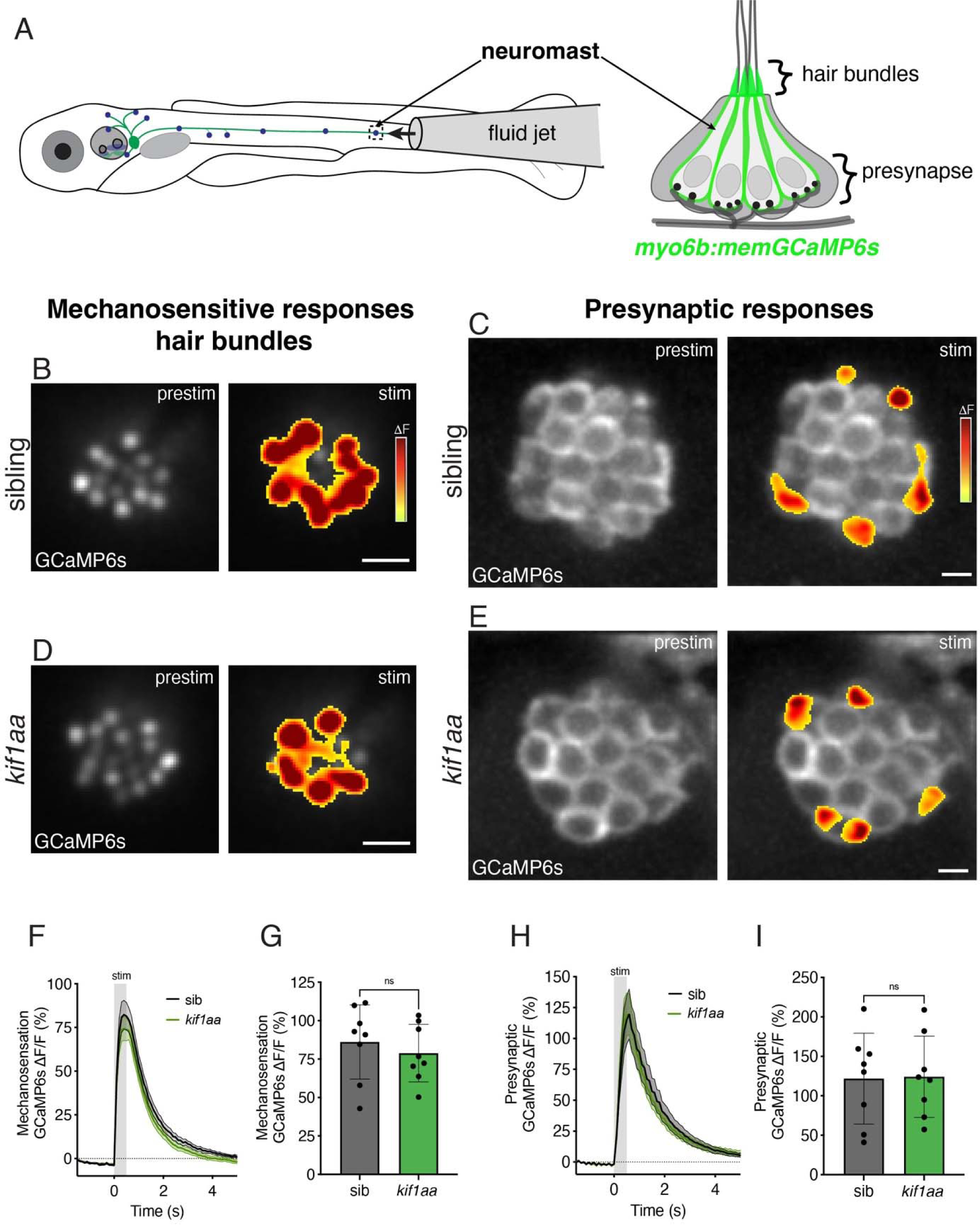
*Kif1aa* mutants have normal mechanosensitive and presynaptic responses. (**A**) Overview of the scheme used to assess evoked calcium responses in lateral-line hair cells. A fluid-jet is used to deliver flow stimuli to lateral-line neuromasts. A membrane-localized GCaMP6s (*myo6b:memGCaMP6s*, green) expressed in hair cells is used to measure fluid-jet evoked calcium signals in apical hair bundles or presynaptic calcium signals at the cell base. (**B-E**) Top-down images show optical planes of memGCaMP6s in neuromast hair bundles (**B,D**) or at the presynapse (**C,E**). Heatmaps show spatial representations of Δ GCaMP signals during evoked mechanosensitive (**B-C**) and presynaptic (**D-E**) activity during a 500-ms stimulation (stim) compared to prestimulus (prestim) in sibling controls and *kif1aa* mutants. (**F-I**) Traces show the average mechanosensitive (**F**) and calcium presynaptic (**H**) calcium responses in sibling control and *kif1aa* mutant hair cells (n = 8 neuromasts). Dot plots show that the average mechanosensitive (**G**) and presynaptic (**I**) calcium responses are similar in sibling control and *kif1aa* mutant hair cells (**G**, control: 86.1 ± 24.2, *kif1aa*: 78.9 ± 18.7, n = 8 control and *kif1aa* neuromasts, unpaired t-test, p = 0.514; **I**, control: 121.7 ± 57.6, *kif1aa*: 124.1 ± 51.5, n = 8 control and *kif1aa* neuromasts, unpaired t-test, p = 0.930, 5 dpf). Scale bar in **B-E** = 5 µm.

To obtain a more sensitive readout of hair-cell mechanosensation we used stimulus-evoked calcium imaging. We stimulated lateral-line hair cells with a fluid jet to evoke activity (Figure 7A) and read out calcium activity using a transgenic line that expresses membrane-localized GCaMP6s specifically in hair cells (*myo6b:memGCaMP6s*) (Lukasz & Kindt, 2018; Zhang *et al*., 2018). This transgenic line can be used to image mechanosensation-dependent calcium signals in apical hair bundles, as well as calcium signals at the base of the cell that are associated with presynaptic-calcium influx (Figure 7A-B,D). We first examined calcium signals in apical hair bundles. In response to a 500-ms stimulus, we observed similar response magnitudes (ΔF/F GCaMP6s) in *kif1aa* mutants compared to sibling controls (Figure 7B,D,F,G, control: 86.1 ± 24.2; *kif1aa*: 78.9 ± 18.7, n = 8 control and *kif1aa* neuromasts, unpaired t-test, p = 0.514). Together our FM 1-43 labeling and calcium imaging experiments indicate that mechanosensation is intact in lateral-line hair cells of *kif1aa* mutants.

After assessing mechanosensation, we used the same transgenic line (*myo6b:memGCaMP6s*) and stimulation approach to examine presynaptic calcium responses at the hair-cell base. In responses to a 500-ms stimulus, we observed similar presynaptic-response magnitudes (ΔF/F GCaMP6s) in *kif1aa* mutants compared to sibling controls (Figure 7C,E,H,I, control: 121.7 ± 57.6; *kif1aa*: 124.1 ± 51.5, n = 8 control and *kif1aa* neuromasts, unpaired t-test, p = 0.930). Overall, our data indicate that both mechanosensitive and presynaptic responses in lateral-line hair cells are largely normal in *kif1aa* mutants.

### Loss of Kif1aa alters spontaneous release and postsynaptic calcium responses

Our initial calcium imaging experiments suggests that presynaptic responses in *kif1aa* mutants are relatively normal (Figure 7C). Because *kif1aa* mutants fail to enrich synaptic vesicles at the presynaptic AZ, we hypothesized that vesicle release could still be impaired despite normal presynaptic-calcium responses. Therefore, we used additional functional approaches to examine the postsynaptic outcome of fewer synaptic vesicles.

Similar to hair cells in other species, zebrafish lateral-line hair cells release glutamate-filled synaptic vesicles at rest. Established work has shown that this release generates spontaneous spikes in neurons of the posterior lateral-line ganglion (pLLg)– neurons that innervate posterior lateral-line neuromasts (Figure 1A, (Trapani & Nicolson, 2011)). We used loose-patch electrophysiology recordings to measure spontaneous spikes in pLLg neurons (Figure 8A). To quantify the rate of spontaneous spiking, we measured the number of spikes per minute over a 5-min window. We observed a marked ∼80 % decrease in the average afferent spike rate in *kif1aa* mutants compared to sibling controls (Figure 8B-C, control: 207.5 ± 148.3, *kif1aa*: 42.03 ± 19.70; n = 8 control and 13 *kif1aa* cells, unpaired t-test, p = 0.0007). This indicates that a reduction in synaptic vesicles at the presynapse in *kif1aa* mutants results in fewer vesicles released at rest.

**Figure 8.**
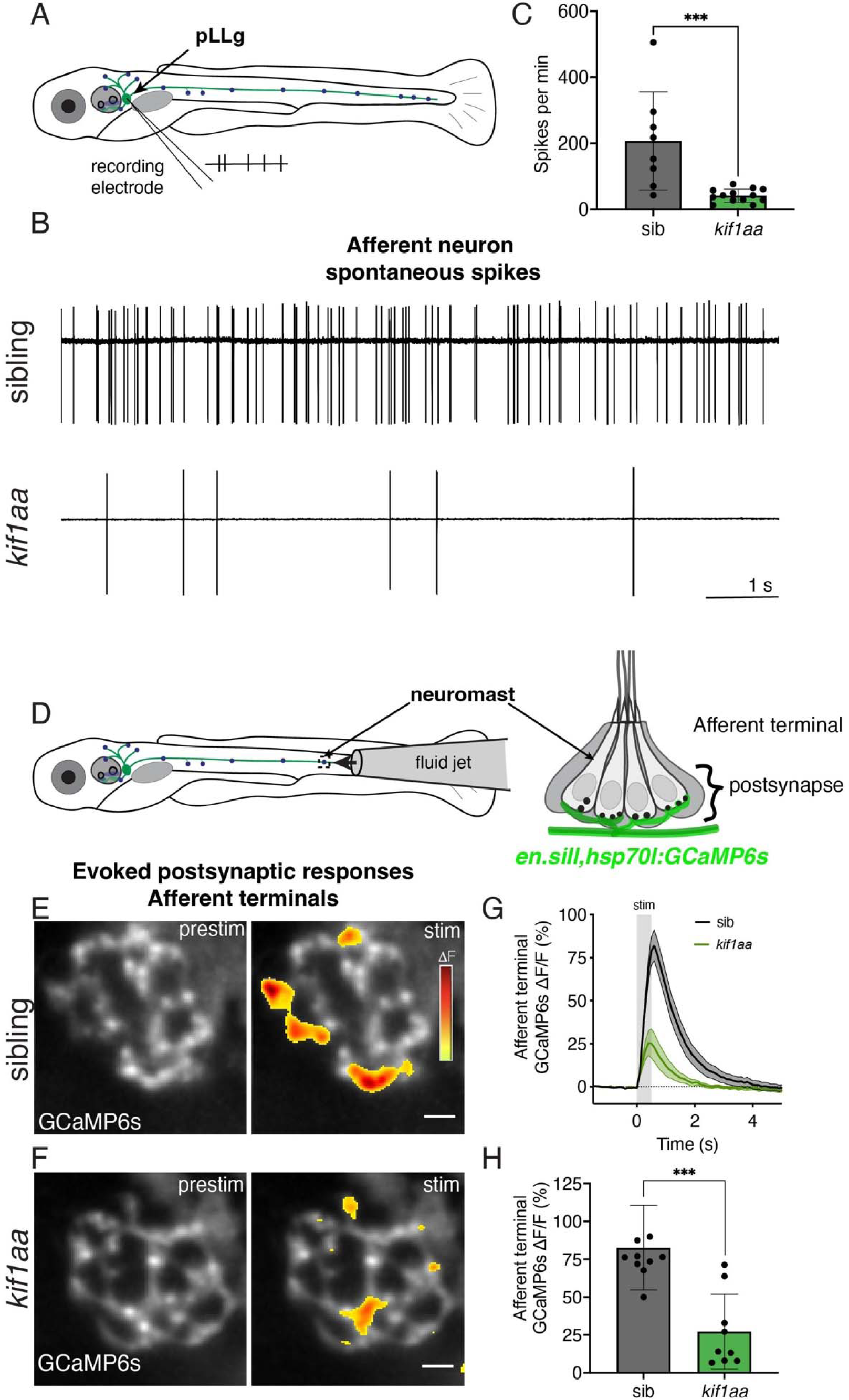
Afferent neurons in *kif1aa* mutants have fewer spontaneous spikes and reduced evoked responses. (**A**) Overview of the scheme used to record spontaneous spiking from afferent cell bodies in the posterior lateral-line ganglion (pLLg). Glass pipettes pulled with long tapers were used to record extracellularly in a loose-patch configuration. (**B**) Representative 10 s traces show spiking in pLLg neurons in sibling control (top) and *kif1aa* mutant (bottom). (**C**) Quantification shows that the average number of spikes per min in *kif1aa* mutants is significantly lower than sibling controls (control: 207.5 ± 148.3, *kif1aa*: 42.03 ± 19.70; n = 8 control and 13 *kif1aa* cells, unpaired t-test, p = 0.0007, 3-6 dpf). (**D**) Overview of the scheme used to assess evoked calcium responses in the afferent terminals beneath lateral-line hair cells. A fluid-jet is used to deliver flow stimuli to lateral-line neuromasts. A transgenic line (*en.sill,hsp70l:GCaMP6s*) expressed in posterior lateral-line afferents is used to measure fluid-jet evoked GCaMP6s calcium signals in afferent terminals beneath neuromasts. (**E-F**) Top-down images show optical planes of GCaMP6s in afferent terminals. Heatmaps show spatial representations of Δ GCaMP signals in afferent terminals during a 500-ms stimulation (stim) compared to prestimulus (prestim) in sibling control and *kif1aa* mutants. (**G**) Traces show the average response in the afferent terminal in sibling control and *kif1aa* mutants (n = 10 control and 9 *kif1aa* neuromasts). (**H**) Dot plot shows that the average response in the afferent terminal is significantly lower in *kif1aa* mutants compared to sibling control (control: 82.6 ± 27.9, *kif1aa*: 27.2 ± 24.7; n = 10 control and 9 *kif1aa* neuromasts, unpaired t-test, p = 0.0003, 4-5 dpf). Scale bar in **E-F** = 5 µm and 1 s in **B**.

To assess evoked postsynaptic activity, we used calcium imaging. For our imaging, we used a transgenic line that expresses GCaMP6s specifically in neurons in the pLLg (*en.sill,hsp70l:GCaMP6s*) (Zhang *et al*., 2018). We stimulated hair cells with a fluid jet and used this line to measure evoked calcium signals in the afferent terminal beneath lateral-line hair cells (Figure 8D-F). In responses to a 500-ms stimulus, we observed that postsynaptic calcium responses (ΔF/F GCaMP6s) in *kif1aa* mutants were dramatically reduced compared to sibling controls (Figure 8E-H, control: 82.6 ± 27.9; *kif1aa*: 27.2 ± 24.7, n = 10 control and 9 *kif1aa* neuromasts, unpaired t-test, p = 0.0003). Together, our postsynaptic electrophysiology and calcium imaging indicate that loss of Kif1aa and fewer synaptic vesicles leads to impaired release of synaptic vesicles.

### Kif1aa is required for normal rheotaxis behavior, but not acoustic startle behavior

After verifying that loss of Kif1aa impaired synapse function in lateral-line hair cells, we assessed what impact this impairment had on behavior. We examined two hair-cell mediated behaviors in zebrafish: the acoustic startle response and rheotaxis. The acoustic startle response is well-characterized zebrafish behavior where a rapid escape reflex occurs in response to an acoustic or vibrational stimulus (Kimmel *et al*., 1974; Granato *et al*., 1996). This behavior relies primarily on hair cells in the posterior macula (saccule), as well as hair cells in the lateral line (Granato *et al*., 1996; Holmgren *et al*., 2021). Rheotaxis, whereby fish orient into an oncoming current and hold their position to avoid being swept downstream, is a multisensory behavior where the lateral-line organ is used to sense changes in water flow (Coombs *et al*., 2020). Loss of lateral-line function in larval zebrafish leads to an inability to station hold (i.e., hold position at the source of oncoming water flow) during bouts of rheotaxis behavior (Newton *et al*., 2023).

To assay the acoustic startle response in our *kif1aa* mutants we used a Zantiks automated behavioral system. We presented larvae in a 12-well plate with acoustic vibrational stimuli of 3 levels of intensity (level 3 highest, level 1 lowest), with 5 trials per intensity level. Each trial was done with a 2-min inter-stimulus interval (ISI) to avoid habituation. Using Zantiks software, larvae were recorded and tracked throughout the experiment (Figure 9-S1A-B). We found that our *kif1aa* mutants were able to respond to a single non-habituating stimulus at the same rate as sibling controls at each stimulus intensity level (Figure 9-S1C, n = 38 control and 13 *kif1aa* larvae, 2-way ANOVA, level 3: p = 0.999, level 2: p = 0.999, level 1: p = 0.994, no stimulus: p = 0.586, 5 dpf). Because we saw no difference in *kif1aa* mutants in our acoustic startle assay, we decided to push this assay further to see if our mutants could maintain responses when presented with repeated or habituating stimuli with short ISIs. For this assay we first administered 3 non-habituating stimuli with a 2-minute ISI, followed by a train of 30 habituating stimuli with a 5-second ISI. We also tested recovery from habituation; here we administered one stimulus at 20 s, 40 s, 1 min, and 2 min each after the last habituating stimulus. Using this paradigm, we found that *kif1aa* mutants habituate at the same rate as sibling controls (Figure 9-S1D, n = 38 control and 25 *kif1aa* larvae, 2-way ANOVA with multiple comparisons, across stimulus and genotype, p = 0.545). In addition, the rate of recovery after habituation was not significantly different between *kif1aa* mutants and sibling controls (Figure 9-S1D, n = 38 control and 25 *kif1aa* larvae, 2-way ANOVA with multiple comparisons, across stimulus and genotype, p = 0.620). Overall, our acoustic startle assays indicate that loss of Kif1aa does not impair this escape reflex.

**Figure 9.**
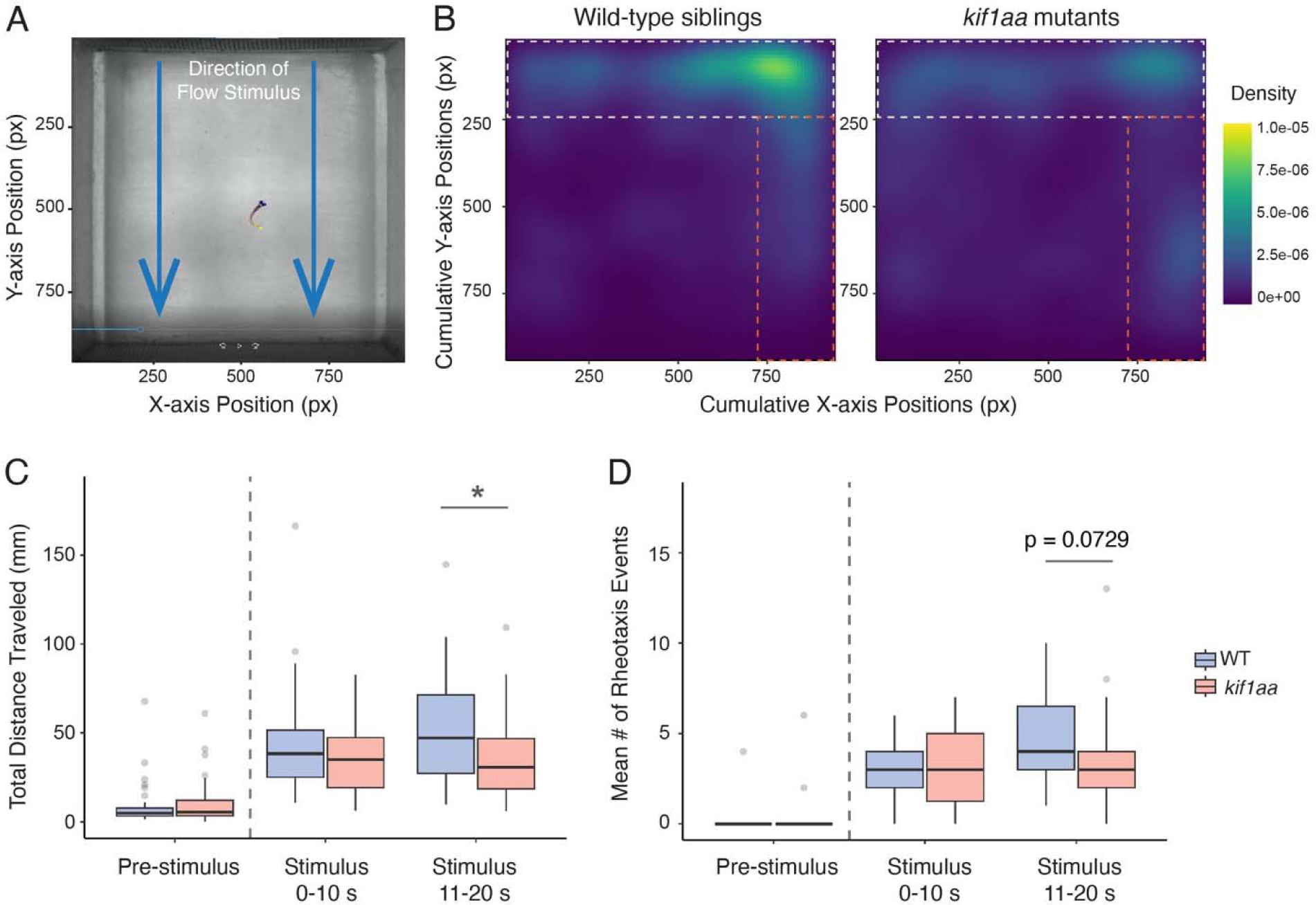
Station holding within flow stimulus is impaired in *kif1aa* mutants. (**A**) Top-down view of the working section of the microflume apparatus. Blue arrows indicate the direction of water flow. A larval fish performing behavior is included for scale. (**B**) Two-dimensional heat maps showing spatial use/cumulative positioning during flow stimulus. Wild-type siblings (**B**) predominantly maintain position in the space at the front of the arena (white dotted lines) in the strongest part of the flow. Cumulative positioning in the source of the flow is reduced for *kif1aa* mutants (white vs. orange dotted lines) indicating impaired ability to station hold. (**C**) Box and whisker plots of total distance traveled during rheotaxis events. Under flow stimulus, the total distance traveled by *kif1aa* mutant larvae was significantly reduced compared to wild-type siblings during the second half (11-20 s) of flow stimulus (control: 50.08 nm; kif1aa: 35.07 nm, adjusted p = 0.017 (11-20 s)). (**D**) Box and whisker plots of the mean number of rheotaxis events. Under flow stimulus, *kif1aa* mutant larvae trended toward fewer rheotaxis events during the last half (11-20 s) of stimulus, though the difference was not significant (control: 4.53 events; kif1aa: 3.70 events, adjusted p = 0.0728). n = 43 wild-type and 30 *kif1aa* mutant larvae were tested.

Our immunostaining demonstrated that synaptic-vesicle enrichment was less affected in hair cells of the inner ear in *kif1aa* mutants (Figure 3-S2). This may allow inner ear hair cells in *kif1aa* mutants to function normally, as the acoustic startle is a fast reflex and may not require sustained synaptic vesicle release. Therefore, we examined rheotaxis behavior, a hair-cell mediated behavior that is reliant on lateral-line hair cells, as synaptic vesicle populations in these hair cells were dramatically impaired in *kif1aa* mutants. To evaluate rheotaxis behavior, we used a custom microflume to present laminar flow to zebrafish at 6 dpf, at a rate of ∼10 mm s^-1^ (Figure 9A; (Baker & Montgomery, 1999; Newton *et al*., 2023)). We recorded the behavior of larvae with an overhead camera 10 s before flow (pre-stimulus) and for 20 s during flow stimulus using infrared (850 nm) illumination to eliminate visual cues. We measured spatial use of the chamber, distance traveled, and rheotaxis events (orienting in flow) during flow. We first we examined the spatial use of the flow chamber, we found that when performing rheotaxis, wild-type siblings were predominantly positioned in the space at the front of the arena, in the region of the highest flow (Figure 9B; white dashed box). In contrast, *kif1aa* mutants cumulatively show less positioning in high-flow regions, indicating a reduced ability to station hold (Figure 9B). During bouts of rheotaxis, we also observed that *kif1aa* mutants accumulated along the side of the arena in a small lateral gradient in the laminar flow field (Figure 9B; orange dashed box). Together, these observations indicate that *kif1aa* mutants have an impaired ability to maintain position during rheotaxis. To determine why *kif1aa* mutants showed a reduced station holding, we evaluated whether there were differences in the metrics of rheotaxis behavior for the first (0-10 s) and last half (10-20 s) of stimulus. When we examined distance traveled during rheotaxis, we found that during the last half of the stimulus, *kif1aa* mutants traveled a reduced total distance compared to wild-type siblings (Figure 9C, 0-10 s, control: 43.33, *kif1aa*: 37.26, p = 0.532; 11-20 s, control: 50.08, *kif1aa*: 35.07, p = 0.0728, n = 43 controls and 30 *kif1aa* larvae). Importantly, we did not observe a difference in total distance traveled before the onset of flow, indicating that motor function is largely intact in *kif1aa* mutants (Figure 9C). Further, we observed a trend toward fewer rheotaxis events in *kif1aa* mutants during the last half of stimulus, though it did not reach significance (Figure 9D, 0-10 s, control: 3.21, *kif1aa*: 3.03, p = 0.532; 11-20 s, control: 4.53, *kif1aa*: 3.70, p = 0.0728, n = 43 controls and 30 *kif1aa* larvae).

Taken together, our acoustic startle and rheotaxis assays demonstrate that while *kif1aa* mutants can respond normally to discrete acoustic stimuli, their ability to effectively station hold during rheotaxis behavior in sustained flow stimuli is impaired. This suggests that a large population of synaptic vesicles is important for behaviors that require sustained neurotransmission.

## Discussion

Our work in zebrafish demonstrates that both Kif1a and an intact microtubule network are essential to localize synaptic vesicles properly at the presynaptic AZ. Our functional assays reveal that hair cells lacking Kif1a release fewer synaptic vesicles, leading to impaired afferent responses. Importantly, deficits in hair-cell vesicle release results in impaired station-holding during rheotaxis–a behavior mediated by the lateral-line sensory system. Altogether, our work highlights a new pathway that ensures an adequate supply of synaptic vesicles reaches the cell base to maintain high rates of release at specialized ribbon synapses.

### Accessory machinery and cargo in transport vesicles

Our work demonstrates that lateral-line hair cells lacking Kif1aa have less enrichment of synaptic markers (Figure 3, Figure 3-S1) and fewer synaptic vesicles at the presynaptic AZ (Figure 5). This finding is consistent with studies in neurons in both mice and *C. elegans* where disruptions in Kif1aa orthologues Kif1a/UNC-104 result in significantly fewer synaptic vesicles at presynapses (Hall & Hedgecock, 1991; Okada *et al*., 1995). In neurons, synaptic-vesicle precursors are made *de novo* in the ER-Golgi network located in the cell soma. After budding from the trans-Golgi, these precursors are carried to presynaptic terminals along axonal microtubules via kinesin motors and mature into synaptic vesicles at the presynapse (Santos *et al*., 2009; Rizzoli, 2014). Precursor vesicles can contain many presynaptic components including: Rab3a, CSP, Vglut, Piccolo, Bassoon, SNARE proteins, Neurexins and calcium channels (reviewed in: (Petzoldt, 2023)). In our *kif1aa* mutants we show that several of these cargos such as Rab3a, CSP, and Vglut3 fail to localize properly at the presynaptic AZ in hair cells (Figures 3, Figure 3-S1). Although we did observe more Ca_V_1.3 channels per presynapse (Figure 6F), we also observed fewer complete synapses (Figure 2D) and fewer Rib b-Ca_V_1.3 paired puncta (Figure 6C) in *kif1aa* mutants. Fewer complete pairings could be due to a requirement for Kif1aa to transport material (ex: Ribeye and Ca_V_1.3) during synapse formation. This is consistent with our previous work that revealed subtle defects in ribbon formation in lateral-line hair cells lacking Kif1aa (Hussain *et al*., 2024). Overall, our work on hair cells suggests that Kif1a plays a conserved role in the transport of synaptic material to the presynaptic AZ of neurons and hair cells.

An important part of transport is not only the kinesin motor moving cargo along microtubules, but also adaptors that link specific cargos to motors. Work in neurons suggests that the Kif1a motor relies on the adaptor protein MAP kinase activating death domain (Madd) to transport synaptic vesicle precursors. Specifically, Madd is thought to provide a linkage between Rab3a present on synaptic-vesicle precursors and Kif1a (Niwa *et al*., 2008; Hummel & Hoogenraad, 2021). In our study, and other work on hair cells, Rab3a has been shown to be a marker of synaptic vesicles in hair cells (Uthaiah & Hudspeth, 2010; Einhorn *et al*., 2012). Along with other synaptic markers, we found that in *kif1aa* mutants, Rab3a immunolabel was not enriched at the hair-cell presynapse (Figure 3-S1E-H). Based on this result and work in neurons, it is possible that Rab3a is part of the link that is required to facilitate the transport of new synaptic-vesicle precursors from the Golgi in hair cells. In the future it will be interesting to examine the localization of various synaptic vesicle markers in *rab3a* mutants or to lesion other transport partners such as Madd to determine whether they are also required to accumulate synaptic vesicles at the presynapse in hair cells.

### Mechanisms to maintain synaptic vesicle populations at the presynapse

We found that fewer synaptic vesicles in *kif1aa* mutants were accompanied by impaired spontaneous and evoked activity in lateral-line afferent neurons (Figure 3, Figure 5, Figure 8). But despite a dramatic reduction in synaptic vesicles and activity, some synaptic vesicles were still present at ribbon synapses and activity was not completely abolished (Figures 5, Figure 8). Based on these results it is likely that there are alternate mechanisms in play that ensure synaptic vesicles reach the presynapse. For example, other kinesin motors could work in tandem with Kif1a to transport synaptic vesicle precursors. In support of this idea, immunolabeling results have shown that another kinesin, Kif3a, colocalizes with ribbons (Michanski *et al*., 2019). Alternatively, it is possible that synaptic-vesicle precursors may simply reach the presynapse through diffusion. Studies on ribbon synapses suggest that synaptic vesicles are able to freely diffuse within the cytosol until affixing to the ribbon (Holt *et al*., 2004; LoGiudice & Matthews, 2009). Based on these diffusion studies, it is possible that synaptic and precursor vesicles in *kif1aa* mutants diffuse from their site of origin near the Golgi until they eventually encounter and bind to a ribbon. In this scenario, although diffusion is less efficient than directed transport, it may allow enrichment of enough synaptic vesicles to explain the residual function at synapses in *kif1aa* mutants.

In many types of neurons, especially those with long axons, it is essential that new synaptic-vesicle precursors are actively transported, as presynaptic terminals are located at considerable distances from the cell soma. In these circumstances, the diffusion of synaptic-vesicle precursors is not practical. In addition to *de novo* synthesis and transport of synaptic material from the Golgi to the presynapse, at both neuronal synapses and ribbon synapses, endocytosis and local synaptic vesicle recycling also function to maintain synapse function. For example, work in hippocampal neurons has demonstrated that robust recycling of synaptic vesicles can maintain stable and consistent synaptic release even during high frequency and sustained stimulation (Gallimore *et al*., 2023). In hair cells, multiple forms of endocytosis have been characterized at ribbon synapses including clathrin-mediated endocytosis and bulk endocytosis; there is also evidence of kiss-and-run fast endocytosis (Neef *et al*., 2014). In the future, it will be important to more carefully explore whether endocytic pathways are disrupted in hair cells lacking Kif1aa.

While there are several pathways in hair cells capable of recycling synaptic material, it is less clear where this recycling occurs. For example, most of the recycling could occur locally at the presynaptic AZ. Synaptic material could also reenter the cell apically and be re-trafficked to the presynapse at the cell base. Additionally, it is possible that both apical and basal locations could be used for recycling synaptic material. Electron micrographs show evidence that both clathrin-mediated endocytosis and bulk endocytosis occur near the presynapse (Siegel & Brownell, 1986; Lenzi *et al*., 2002; Kamin *et al*., 2010). In addition, studies using FM 1-43 uptake to measure endocytosis in hair cells have shown that endocytosis may also occur more apically and undergo recycling in the Golgi (Griesinger *et al*., 2002, 2005). Endocytosis in hair cells has also been studied using the membrane tracer mCLING (Revelo *et al*., 2014). Work using mCLING revealed evidence that synaptic material is recycled apically, as well as basally at the presynapse via local recycling of synaptic vesicles and bulk endocytosis. Overall, it is likely that *de novo* synthesis and transport of synaptic material from the Golgi, bulk endocytosis, and local synaptic vesicle recycling all work together to supply synaptic vesicles for release at ribbon synapses. In the future it would be interesting to create transgenic zebrafish that label synaptic vesicles in hair cells. *In vivo* imaging of this transgenic line could be used to understand how synaptic vesicles are transported and recycled in living hair cells–and how these processes are perturbed in *kif1aa* mutants.

### Impact of Kif1a loss on zebrafish behavior and in human patients

In humans mutations in the *KIF1A* gene lead to a rare inherited condition collectively known as KIF1A-associated neurological diseases (KANDs) (Nair *et al*., 2023). Because *KIF1A* is expressed broadly in the nervous system, there is extensive damage to neurons in the brain and spinal cord in KAND patients. Further, while optic nerve atrophy and vision impairment are also a part of KAND, hearing loss is a less well-studied symptom of this disease (Lee *et al*., 2015; Montenegro-Garreaud *et al*., 2020; Pennings *et al*., 2020). Our work suggests that in addition to neurons, KIF1a may also play an important role in sensory hair cells. Our study was able to highlight this role more clearly because zebrafish have two ohnologues of mammalian KIF1a, Kif1aa and Kif1ab (Figure 1I). Based on published scRNAseq data (Sur *et al*., 2023), in our *kif1aa* zebrafish mutants it is likely that Kif1ab compensates for loss of Kif1aa in most neurons. This allowed us to focus on phenotypes relevant to hair cells in the lateral line, which only express *kif1aa* (Figure 1F-H). Behaviorally, our *kif1aa* mutants have normal acoustic startle responses (Figure 9-S1). Normal startle behavior could be due to contributions from hair cells in the zebrafish inner ear which are less affected in our *kif1aa* mutants (Figure 3-S2). Alternatively, it is possible that fewer synaptic vesicles are needed for this reflexive behavior. Importantly, our rheotaxis experiments demonstrated that *kif1aa* mutants are unable to maintain position in flow (Figure 9B). Further, the deficits in rheotaxis behavior occurred during the last 10 s of stimulus (Figure 9C, D), suggesting that the inability to maintain synaptic vesicles at hair-cell presynapses may underlie these deficits. The continuous flow applied during our rheotaxis experiments demands a near constant supply of synaptic vesicles, specifically from lateral-line hair cells. In future studies, it will be important to rescue Kif1a in hair cells to ensure that our behavioral phenotypes are due to defects that arise specifically in hair cells.

Overall, our work suggests that inner ear deficits may contribute to phenotypes in human KAND patients. For example, KAND is associated with cerebellar ataxia, and although there is evidence of cerebellar atrophy in KAND patients, it is possible that there are also contributions from the inner ear, particularly the vestibular system where hair cells must maintain sustained release of synaptic vesicles for proper balance (Paprocka *et al*., 2023). In the future it will be important to assess KAND patients carefully for hearing and balance deficits. In addition, our *kif1aa* zebrafish could be used to understand the role KIF1a plays in hair cells or to assess pathogenic variants in KIF1a.

Our current study highlights how an understudied mechanism–Kif1a-based transport– functions to enrich synaptic vesicles at ribbon synapses in hair cells. In the future it will be important to understand how this transport works alongside recycling and endocytic pathways to supply the near constant demand for synaptic vesicles at ribbon synapses. A powerful way to study synaptic vesicle dynamics is by using live imaging and sensitive readouts of release, such as afferent recordings. The zebrafish lateral line provides an excellent model to apply these approaches in future studies.

## Methods

### Zebrafish husbandry

Zebrafish (*Danio rerio*) lines were maintained at the National Institutes of Health (NIH) under animal study protocol #1362-13. Zebrafish larvae (0-7 dpf) were raised in incubators at 28°C in E3 embryo medium (5 mM NaCl, 0.17 mM KCl, 0.33LmM CaCl_2_, and 0.33 mM MgSO_4_, buffered in HEPES, pH 7.2) with a 14hr:10hr light:dark cycle. Lines were maintained in a Tu or TL background.

For rheotaxis experiments embryos were shipped at 1 dpf from NIH to Washington University-St. Louis and maintained under IACUC protocol #23-0078, and subsequently, raised in incubators at 28°C in embryo media2 (15 mM NaCl, 0.5 mM KCl, 1 mMCaCl_2_, 1 mM MgSO_4_, 0.15 mM KH_2_PO_4_, 0.042 mM Na_2_HPO_4_, 0.714 mM NaHCO_3_) with a 14hr:10hr light:dark cycle. After 4 dpf, larvae were raised in 100-200 ml of EM in 250 ml plastic beakers and fed rotifers daily.

All experiments were performed on larvae at 3-6 days post fertilization (dpf). Neuromasts L1-L4 were used for all analyses, except the Ca_V_1.3 immunolabel where L1-L4, D1 and O1 were examined, and our TEM analysis where cranial neuromasts M1, along with the POs, and SOs were examined. Larvae were chosen at random and used at an age where sex determination is not possible. The previously described mutant and transgenic lines were used in this study: *kif1aa^idc24^*, *Tg(myo6b:memGCaMP6s)^idc1Tg^, Tg(en.sill,hsp70l:GCaMP6s)^idc8Tg^, Tg(myo6b:Cr.ChR2-EYFP)^ahc1Tg^, Tg(myo6b:ctbp2a-TagRFP)^idc11Tg^* (also referred to as: *Tg(myo6b:rib a-TagRFP)^idc11Tg^*) and *Tg(myo6b:YFP-Hsa.TUBA)^idc16Tg^* (Monesson-Olson *et al*., 2014; Zhang *et al*., 2018; Wong *et al*., 2019; Ohta *et al*., 2020; Hussain *et al*., 2024)

The *kif1aa* germline mutant has been previously described (Hussain *et al*., 2024). This Crispr-Cas9 mutant has a complex INDEL that leads to a disruption in Kif1aa at amino acid 166, in exon 6, within the kinesin motor domain (Figure 1I). This INDEL also disrupt a Bsl1 restriction site in exon 6. Genotyping was done using standard PCR followed by a Bsl1 restriction enzyme digest. *Kif1aa* genotyping primers used were as follows: *kif1aa*_FWD 5’-AACACCAAGCTGACCAGTGC-3’ and *kif1aa*_REV 5’-TGCGGTCCTAGGCTTACAAT-3’. *Kif1aa* mutants were compared to either heterozygous and wild-type siblings (sibling controls) or to wild-type siblings as stated in the text.

### Nocodazole treatment

To destabilize microtubules, *Tg(myo6b:YFP-Hsa.TUBA)^idc16Tg^* larvae at 5 dpf were incubated in 250 nM nocodazole (Sigma-Aldrich, SML1665) for 2 hours. Nocodazole was diluted in E3 media for a final concentration of 250 nM nocodazole and 0.1 % DMSO. After effective nocodazole treatment *YFP-Hsa.TUBA* labeling was visually disrupted. For controls, larvae were incubated in media containing 0.1 % DMSO. After 2 hours, larvae were fixed for immunohistochemistry or prepared for LysoTracker labeling (see below).

### Lysotracker labeling, and imaging

After 2 hours of nocodazole treatment, 100 nM LysoTracker Red DND-99 (ThermoFisher, L7528) was added to the media for 15 min. Larvae were then embedded in 1 % low melt agarose prepared in E3 media containing 0.03 % tricaine (Sigma-Aldrich, A5040, ethyl 3-aminobenzoate methanesulfonate salt), 100 nM LysoTracker and either 250 nM nocodazole or 0.1 % DMSO. A similar labeling approach was used for Lysotracker labeling in *kif1aa* mutants. LysoTracker Red DND-99 was used to label *Tg(myo6b:YFP-Hsa.TUBA)^idc16Tg^* larvae while LysoTracker Green DND-26 (ThermoFisher, L7526) was used to label *Tg(myo6b:rib a-TagRFP)^idc11Tg^* larvae. Transgenic larvae were incubated in 100 nM LysoTracker dye in E3 media for 15 min and mounted in 1 % low melt agarose prepared in E3 media containing 0.03 % tricaine and 100 nM Lysotracker dye.

To image LysoTracker label, samples were imaged live on a Nikon A1R upright confocal microscope using a 60x 1 NA water objective lens. Denoised images were acquired using NIS Elements AR 5.20.02 with an 0.425 µm z-interval. Z-stacks of whole or partial neuromasts were acquired in a top-down configuration using 488 and 561 nm lasers.

### FM 1-43 labeling, imaging and analysis

To assay mechanotransduction using FM 1-43, larvae at 5 dpf were incubated in 3 µM FM 1-43 (Thermofisher, T3163) in E3 media for 35 s. Larvae were then washed 3× in E3 and mounted in 1 % low melt agarose prepared in E3 media with 0.03 % tricaine. After mounting larvae were imaged on a Nikon A1R upright microscope using a 60x 1 NA water objective lens. Denoised images were acquired using NIS Elements AR 5.20.02 with an 0.425 µm z-interval. Z-stacks of whole neuromasts were acquired in a top-down configuration using 488 and 561 nm lasers. Z-stacks were further processed in FIJI. Z-stacks were max-projected and thresholded to obtain the mean intensity of the FM 1-43 label per neuromast.

### Immunohistochemistry

Immunohistochemistry was performed on whole larvae. Zebrafish larvae were fixed with 4 % paraformaldehyde in PBS for 3-4Lhrs at 4 °C. After fixation samples were washed 5L×L5Lmin in PBSL+L0.01 % Tween (PBST), followed by a 5-min wash in H_2_O. Larvae were then permeabilized with ice cold acetone (at -20 °C) for 5 min. Larvae were then washed again in H_2_O for 5 min, followed by a 5L×L5-min washes in PBST, and then blocked overnight at 4 °C with PBST containing 2 % goat serum, 2 % fish skin gelatin and 1 % bovine serum albumin (BSA). Primary antibodies were diluted in PBST containing 1 % BSA. Larvae were incubated in primary antibodies overnight at 4L°C. After 5L×L5Lmin washes in PBST to remove the primary antibodies, larvae were incubated in diluted secondary antibodies (1:1000) in PBST containing 1 % BSA for 3Lhrs at room temperature. After 5L×L5Lmin washes in PBST to remove the secondary antibodies, larvae were rinsed in H_2_O and mounted in ProLong Gold Antifade (ThermoFisher, P36930).

#### Primary antibody list

| Antigen | Species;<br>subtype | Concentration | Catalog or reference |
| --- | --- | --- | --- |
| Pan-Maguk | Mouse; IgG1 | 1:500 | Millipore MABN7 |
| Myo7A | Rabbit | 1:1000 | Proteus 25-6790 |
| Vglut3 C-term | Rabbit | 1:1000 | (Obholzer *et al.*, 2008) |
| Ca<sub>v</sub>1.3 | Rabbit | 1:500 | (Sheets *et al.*, 2011) |
| Ribeye b | Mouse; IgG2a | 1:10,000 | (Sheets *et al.*, 2011) |
| Pan-CTBP | Mouse; IgG2a | 1:1000 | Santa Cruz 55502 |
| Rab3a | Mouse; IgG1 | 1:1000 | Synaptic System 107 011 |
| Acetylated<br>Tubulin | Mouse; IgG1 | 1:5000 | Sigma T7451 |
| CSP | Rabbit | 1:1000 | Sigma AB1576 |
| GFP | Chicken | 1:1000 | Thermofisher A10262 |

#### Secondary antibody list

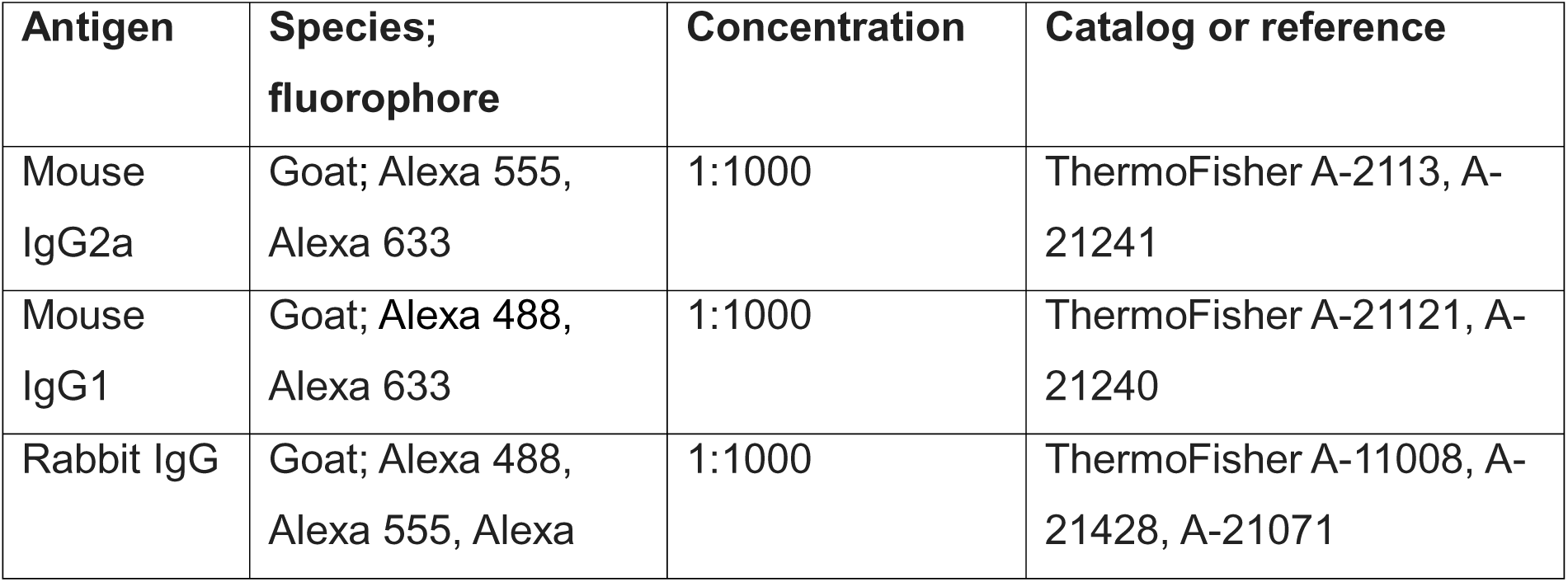
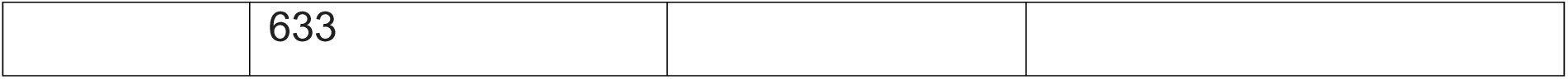

| Antigen | Species;<br>fluorophore | Concentration | Catalog or reference |
| --- | --- | --- | --- |
| Mouse<br>IgG2a | Goat; Alexa 555,<br>Alexa 633 | 1:1000 | ThermoFisher A-2113, A-21241 |
| Mouse<br>IgG1 | Goat; Alexa 488,<br>Alexa 633 | 1:1000 | ThermoFisher A-21121, A-21240 |
| Rabbit IgG | Goat; Alexa 488,<br>Alexa 555, Alexa | 1:1000 | ThermoFisher A-11008, A-21428, A-21071 |

### RNA-FISH to detect *kif1aa* and *kif1ab* mRNA in hair cells

To detect mRNA for *kif1aa and kif1ab* we followed the Molecular Instrument RNA-FISH Zebrafish protocol, Revision Number 10 (https://files.molecularinstruments.com/MI-Protocol-RNAFISH-Zebrafish-Rev10.pdf), with a few minor changes to the preparation of fixed whole-mount larvae as follows. For our dehydration steps we dehydrated using the following methanol series: 25, 50, 75, 100, 100 % methanol, with 5 min for each step in the series. To permeabilize we treated larvae with 10 µg/mL proteinase K for 20 min. RNA-FISH probes were designed to bind after the motor domain of zebrafish *kif1aa and kif1ab* (Figure 1I, Molecular Instrument Probe lot # RTD364, RTD365, using B2 and B3 amplifiers respectively). *Tg(myo6b:Cr.ChR2-EYFP)^ahc1Tg^* larvae were labeled using these probes; the strong EYFP label is retained after RNA-FISH and allows for delineation of hair cells within the whole-mount larvae. After the RNA-FISH protocol the larvae were mounted in ProLong Gold Antifade (ThermoFisher, P36930).

### Confocal imaging of RNA-FISH and immunolabels

Fixed immunostained and RNA-FISH samples were imaged on an inverted LSM 780 laser-scanning confocal microscope with an Airyscan attachment using Zen Blue 3.4 (Carl Zeiss) and an 63x 1.4 NA Plan Apo oil immersion objective lens. Neuromast and inner ear z-stacks were acquired every 0.15 µm. The Airyscan z-stacks were auto processed in 3D. Experiments were imaged with the same acquisition settings to maintain consistency between comparisons. For presentation in figures, images were further processed using Fiji.

### Quantification of RNA-FISH and synaptic components

Z-stack image acquisitions from zebrafish confocal images were processed in FIJI (Schindelin *et al*., 2012). Researchers were blinded to genotype during analyses. Hair-cell numbers were counted manually based on Myo7a, Cr.ChR2-EYFP, YFP-Hsa.TUBA or Acetylated tubulin label. Each channel was background subtracted using the rolling-ball radius method. Then each z-stack was max-intensity projected. A mask was generated by manually outlining the region or interest in the reference channel (ex: hair cells via Myo7a, Acetylated tubulin or *Tg(myo6b:Cr.ChR2-EYFP)*). This mask was then applied to the z-projection of each synaptic component or RNA-FISH channel.

We then used automated quantification to quantify puncta using a customized Fiji-based macro previously described (Jukic *et al*., 2024). In this macro, each masked image was thresholded using an adaptive thresholding plugin by <u>Qingzong TSENG</u> (https://sites.google.com/site/qingzongtseng/adaptivethreshold) to generate a binary image of the puncta (presynaptic, postsynaptic, Ca_V_1.3 cluster or RNA-FISH puncta). Individual synaptic or RNA-FISH puncta were then segmented using the particles analysis function in Fiji. A watershed was applied to the particle analysis result to break apart overlapping puncta. After the watershed, the particle analysis was rerun with size and circularity thresholds to generate ROIs and measurements of each punctum. For particle analysis, the minimum size thresholds of 0.025 μm^2^ (Rib b, Ca_V_1.3, *kif1aa* and *kif1ab* RNA-FISH particles) and 0.04 μm^2^ (Maguk) were applied. A circularity factor of 0.1 was also used for the particle analysis. The new ROIs were applied to the original z-projection to get the average intensity and area of each punctum.

To identify paired synaptic components, images were further processed. Here, the overlap and proximity of ROIs from different channels (ex: pre- and post-synaptic puncta) were calculated. ROIs with positive overlap or ROIs within 2 pixels were counted as paired components. The ROIs and synaptic component measurements (average intensity, area) and pairing results were then saved as Fiji ROIs, jpg images, and csv files.

### Synaptic vesicle quantification of live and fixed labels

Z-stack image acquisitions from live or fixed confocal images were processed and quantified in FIJI. A minimum of 3-6 hair cells per neuromast analyzed. Hair cells with a clear side view were used for base to apex analyses. For base to apex, two rectangular ROIs with an area of 2.25 x 0.65 µm (L x W), were placed above and below the nucleus of each hair cell, in 1 z-slice. Nuclei were identified based on Acetylated tubulin or YFP-Hsa.TUBA labeling. The mean values were measured in all ROIs. The ratio of the base to apex was measured per hair cell to assess the enrichment of synaptic vesicles. Ratio values for hair cells were then averaged for a single neuromast base to apex fluorescence value. To quantify Lysotracker fluorescence at ribbons, a circular ROI was drawn around the ribbon indicated by Rib a-mCherry fluorescence, and the mean value was measured in FIJI.

### Startle behavior

A Zantiks MWP behavioral system was used to assess acoustic startle responses in larvae at 5 dpf as previously described (Giese *et al*., 2023; Jukic *et al*., 2024). The Zantiks system tracks and monitors behavioral responses using an infrared camera at 30 frames per second. During the tracking and stimulation, a Cisco router connected to the Zantiks system was used to relay *x*, *y* coordinates of each larva in every frame. A 12-well plate was used for behavioral analyses. Each well was filled with E3 and 1 larva. All fish were acclimated in the plate within the Zantiks chamber in the dark for 15 min before each test. A vibrational stimulus that triggered a maximal proportion of animals startling in control animals without any tracking artifacts (due to the vibration), was used for our strongest stimuli. We used 4 different levels of intensity (1-4, increasing in intensity), with level 4 as the highest intensity stimulus. For our initial startle assay, each larva was presented with stimuli from intensity levels 1-3, 5 times, with 100 s between trials. For each animal, the proportion of startle responses out of the 5 trials was plotted. For our habituation and recovery assay, a non-habituating stimulus, followed by a habituating stimulus train, and lastly recovery stimulus was performed as previously described (Marsden & Granato, 2015). Our non-habituating stimulus was presented 3 times (intensity level 4), with 100 s between trials. This was followed by a habituating train of 30 stimuli (same stimulus intensity), presented with 5 s in between each stimulus. We then presented recovery stimuli once each at 20 s, 40 s, 1 min, and 2 min after the last stimulus in the habituating train. The proportion of startle responses out of the initial 3 non-habituating stimuli and the proportion of responses at each habituation block and recovery stimulus were plotted. To qualify as a startle response, a distance above 4 pixels or ∼1.9 mm was required within 2 frames after stimulus onset. Larvae were excluded from our analysis if no tracking data was recorded. Startle behavior was performed on at least three independent days.

### Rheotaxis behavior

A custom microflume (previously described in: (Newton *et al*., 2023)) was used as the experimental apparatus for rheotaxis behavior. Laminar water flow of a constant velocity was provided by a 6V bow thruster motor (#108-01, Raboesch) inserted into the flume. An Arduino (UNO R3, Osepp) was programmed with custom scripts to coordinate the timing of the flow and video recording. An array of 196 LEDs emitting infrared light (850 nm) provided illumination for video capture through a layer of diffusion material (several Kimwipes sealed in plastic) and the translucent bottom of the flume. A monochromatic high-speed camera (SC1 without IR filter, Edgertronic.com) with a 60 mm manual focus macro lens (Nikon) was used to record behavioral trials at 60 fps. The flume was filled with EM media (28 °C) and the arena placed within the flume. Due to the heat generated by the IR lights, the temperature was monitored and miniature ice packs (2 × 2 cm; −20 °C) were used to maintain a consistent temperature range of 27–29 °C. For each rheotaxis behavior test, individual 6 dpf larval zebrafish were transferred by pipette to the arena within the flume and their swimming activity was monitored for ∼10 s to ensure that it exhibited burst-and-glide swimming behavior; larva that did not exhibit normal swimming behavior during the pre-trial period were not included in the analysis. Each larva was genotyped after behavior acquisition and analysis. A total of 43 wild-type siblings and 30 *kif1aa* mutant larvae were analyzed. Rheotaxis behavior was assessed blind, prior to genotyping. Data was collected from 6 experimental sessions on separate days.

Larval fish were tracked using DeepLabCut as previously described (Newton *et al*., 2023). In brief, video files were downsampled to 1000 x 1000 px and cropped. A previously created and trained single animal maDLC project was used to annotate seven unique body parts (left and right eyes, swim bladder, four points along the tail) on each larva. Videos were analyzed with the maDLC project, and the detections were assembled into tracklets using the box method. The original videos were overlaid with the newly labeled body parts to check for trackelet accuracy. Misaligned tracklets were manually adjusted. Rheotaxis behavior was annotated and analyzed using a previously created custom Python feature extraction script *(SimBA)* that defined positive rheotaxis events as when the larvae swam into the oncoming water flow at an angle of 0 degrees ± 45 degrees for at least 100 ms. Videos processed through DeepLabCut analysis were converted to AVI format using the SimBA video editor function and imported into SimBA as previously described.

### Calcium imaging and electrophysiology

Larvae for electrophysiology recordings were either in a *Tg(myo6b:Cr.ChR2-EYFP)* transgenic background or a nontransgenic background. For calcium imaging either *Tg(myo6b:memGCaMP6s)^idc1Tg^* or *Tg(en.sill,hsp70l:GCaMP6s)^idc8Tg^* transgenic larvae were used. To prepare larvae for calcium imaging and electrophysiology, 3-6 dpf larvae were anesthetized in 0.03 % Tricaine-S (SYNCAINE/MS-222, Syndel), pinned to a Sylgard-filled perfusion chamber at the head and tail. Then larvae were paralyzed by injection of 125 µM α-bungarotoxin (Tocris, 2133) into the heart cavity, as previously described (Lukasz & Kindt, 2018). Larvae were then rinsed once in E3 embryo media to remove the tricaine. Next, larvae were rinsed three times with extracellular imaging solution (in mM: 140 NaCl, 2 KCl, 2 CaCl_2_, 1 MgCl_2_, and 10 HEPES, pH 7.3, OSM 310±10) and allowed to recover prior to calcium imaging or electrophysiology. Researchers were blind to genotype during the acquisition.

Calcium responses in the hair cells and afferent process were acquired on a Swept-field confocal system built on a Nikon FN1 upright microscope (Bruker) with a 60x 1.0 NA CFI Fluor water-immersion objective. The microscope was equipped with a Rolera EM-C2 EMCCD camera (QImaging), controlled using Prairie view 5.4 (Bruker). GCaMP6s was excited using a 488 nm solid state laser. We used a dual band-pass 488/561Lnm filter set (59904-ET, Chroma). For evoked measurements, stimulation was achieved by a fluid jet, which consisted of a pressure clamp (HSPC-1, ALA Scientific) and a glass pipette (pulled and broken to achieve an inner diameter of ∼50 µm) filled with extracellular imaging solution. A 2-s or 500-ms pulse of positive or negative pressure was used to deflect the hair bundles of mechanosensitive hair cells along the anterior-posterior axis of the fish. For GCaMP6s imaging in hair bundles or at the presynapse the *Tg(myo6b:memGCaMP6s)^idc1Tg^*line was used. For GCaMP6s imaging in the afferent terminal beneath lateral line hair cell, the *Tg(en.sill,hsp70l:GCaMP6s)^idc8Tg^* line was used. GCaMP6s measurements were performed on larvae at 4 and 5 dpf. Each neuromast (L1-L4) was stimulated four times with an inter-stimulus interval of ∼2 min. To acquire GCaMP6s evoked responses, 5 z-slices (0.5 µm step for mechanosensation, 1.5 µm step for presynaptic and 2.0 µm step for the afferent process) were collected per timepoint for 80 timepoints at a frame rate of 20 ms for a total of ∼100 ms per z-stack and a total acquisition time of ∼8 sec. Stimulation began at timepoint 31; timing of the stimulus was triggered by an outgoing voltage signal from Prairie view.

GCaMP6s z-stacks were average projected, registered, and spatially smoothed with a Gaussian filter (size = 3, sigma = 2) in custom-written MatLab software as described previously (Zhang *et al*., 2018). The first 10 timepoints (∼1 sec) were removed to reduce the effect of initial photobleaching. Registered average projections analyzed in Fiji to make intensity measurements using the Time Series Analyzer V3 plugin. Here circular ROIs were placed on hair bundles or synaptic sites; average intensity measurements over time were measured for each ROI, as described previously (Lukasz & Kindt, 2018). GCaMP6s data was excluded in the case of excessive motion artifacts. Presynaptic responses were defined as >20% ΔF/F0. Hair-bundle responses were defined as >20% ΔF/F0. Postsynaptic responses were defined as >5% ΔF/F0 and a minimum duration of 500 ms. Calcium imaging data then plotted in Prism 10 (Graphpad). The first 20 timepoints were averaged to generate an F0 value, and all responses were calculated as ΔF/F0. The Area under the curve (AUC) function of Prism was used to determine the peak value for each response. Responses presented in figures represent average responses within a neuromast. The max ΔF/F0 was compared between sibling and *kif1aa* mutants.

Extracellular postsynaptic current recordings from afferent cell bodies of the posterior lateral-line ganglion (pLLg) of zebrafish at 3-6 dpf were performed as previously described (Trapani & Nicolson, 2011; Lukasz *et al*., 2022). Briefly, borosilicate glass pipettes (Sutter Instruments, BF150-86-10 glass with filament) were pulled with a long taper, with resistances between 5 and 15 MΩ. The pLLg was visualized using an Olympus BX51WI fixed stage microscope equipped with a LumPlanFl/IR 60x 1.4 NA water dipping objective (N2667800, Olympus). An Axopatch 200B amplifier, a Digidata 1400A data acquisition system, and pClamp 10 software (Molecular Devices, LLC) were used to collect signals. To record spontaneous extracellular currents, afferent cell bodies were recorded using a loose-patch configuration with seal resistances ranging from 20 to 80 MΩ. Recordings were done in voltage-clamp mode, and signals were sampled at 50 μs/point and filtered at 1 kHz. The number of spontaneous events from one neuron per min was quantified from a 5-min recording window using Igor Pro (Wavemetrics).

### Transmission electron microscopy

Larvae were genotyped at 2 dpf using a larval fin clip method to identify *kif1aa* and wild-type siblings to prepare for TEM (Wilkinson *et al*., 2013). Larval tail DNA was genotyped as described above. At 5 dpf *kif1aa* and wild-type siblings were fixed in freshly prepared solution containing 1.6 % paraformaldehyde and 2.5 % glutaraldehyde in 0.1LM cacodylate buffer supplemented with 3.4% sucrose and 2 µM CaCl_2_ for 2 h at room temperature, followed by a 24-h incubation at 4 °C in a fresh portion of the same fixative. After fixation, larvae were washed with 0.1LM cacodylate buffer with supplements, and post-fixed in 1 % osmium tetroxide for 30 min, and then washed with distilled water. Larvae were, dehydrated in 30 – 100 % ethanol series, which included overnight incubation in 70 % ethanol containing 2% uranyl acetate, and in propylene oxide, and then embedded in Epon. Transverse serial sections (60-70Lnm thin sections) were used to section through neuromasts. Sections were placed on single slot grids coated with carbon and formvar, and then sections were stained with uranyl acetate and lead citrate. All reagents and supplies for TEM were from (Electron Microscopy Sciences). Samples were imaged on a JEOL JEM-2100 electron microscope (JEOL Inc.). Whenever possible, serial sections were used to restrict our analysis to central sections of ribbons adjacent to the plasma membrane and a well-defined postsynaptic density.

To quantify ribbon area, ROIs were drawn in FIJI outlining the electron-dense ribbon, excluding the filamentous “halo” surrounding the ribbon. Vesicles with a diameter of 30–50 nm and adjacent (within 60 nm of the ribbon) to the “halo” were counted as tethered vesicles. Readily releasable vesicles were defined as tethered vesicles between the ribbon and the plasma membrane. To quantify reserve vesicles, we counted vesicles that were not tethered to the ribbon but were within 200 nm of the edge of the ribbon. All distances and perimeters were measured in FIJI.

### Statistics

All data shown are mean and standard deviation unless stated otherwise. All replicates were biological-distinct animals and cells. Wild-type zebrafish animals were selected at random for drug treatments. *Kif1aa* mutants were compared to sibling controls which were either wild-type siblings or a combination of wild-type and or *kif1aa* heterozygous siblings obtained from the same clutch. All experiments were performed blind to genotype. In all datasets dot plots represent the ‘n’. N represents either the number of neuromasts, hair cells, synapses or puncta as stated in the legends. All experiments were replicated on multiple independent days. For our zebrafish experiments a minimum of 3 animals and 8 neuromasts were examined. An exception is our TEM experiments where we examined ribbon synapses in 7 neuromasts from 4 wild-type siblings and 4 neuromasts from 3 *kif1aa* mutants. Sample sizes were selected to avoid Type 2 error. Statistical analyses were performed using Prism 10 software (GraphPad) or in R for our rheotaxis assays. A D’Agostino-Pearson normality test was used to test for normal distributions. To test for statistical significance between two samples with normal distributions, an unpaired t-test was used. For acoustic startle assays, a 2-way ANOVA with multiple comparisons was used. A Sidak test was used to correct for multiple comparisons. For rheotaxis behavior: data wrangling, cleaning, and figure generation were performed with R and performed as previously described (Newton *et al*., 2022). The mean duration, number, total distance traveled, and mean latency to the onset of rheotaxis events were calculated for 10 s bins (pre-stimulus, first 10 s of stimulus, and last 10 s of stimulus) using SimBA and analyzed in R using generalized linear mixed models (GLMM) and post hoc t-tests (Satterthwaite method). Type III ANOVAs provided significance values for the fixed effects of the GLMMs (see Supplementary Tables 1,2). A p-value less than 0.05 was considered significant. All error bars and ± are standard deviation, unless stated otherwise.

## Author contributions

KK and SD did the immunohistochemistry, along with the imaging and analysis of fixed samples. KP did the *kif1aa* and *kif1ab* RNAFISH. ZL analyzed the HCR, synapse and Ca_V_1.3 immunolabels. SD did the electrophysiology and KK did the calcium imaging. YS prepared and imaged samples for TEM. SD did the startle assays. KN, DL and LS did the rheotaxis assay and analysis. LS made the rheotaxis figure. KK and SD made figures and wrote the manuscript. SD, KP, ZL, JS, LS and KK edited and proofed the manuscript.

## Acknowledgements and funding

We thank Drs. Hiu-tung Wong, Catherine Drerup, Elizabeth Cebul and Doris Wu for their thoughtful comments on our manuscript. This work was supported by National Institute on Deafness and Other Communication Disorders (NIDCD) Intramural Research Program Grants 1ZIADC000085-01 (KK), ZICDC000081 (YS), NIDCD Extramural Research Program Grants R01DC016066 (LS), and “Development of Clinician/Researchers in Academic ENT” training grant, award number T32DC000022 (DL). The content is solely the responsibility of the authors and does not necessarily represent the official view of the funding sources.

## Data and code accessibility

Raw data and code reported in this study have been deposited and are publicly available as of the date of publication. The DOIs are {insert here upon acceptance}. For the rheotaxis assay, all R-analysis scripts, details of apparatus construction, and protocols are provided online in the Open-Source Framework repository, https://osf.io/rvyfz/.

All data reported in this paper will be shared by the lead contact upon request. Any additional information required to reanalyze the data reported in this paper is available from the lead contact upon request.

## References

Baker CF & Montgomery JC (1999). The sensory basis of rheotaxis in the blind Mexican cave fish, Astyanax fasciatus. J Comp Physiol A 184, 519–527.

Bhandiwad AA, Zeddies DG, Raible DW, Rubel EW & Sisneros JA (2013). Auditory sensitivity of larval zebrafish (Danio rerio) measured using a behavioral prepulse inhibition assay. J Exp Biol 216, 3504–3513.

Brandt A, Khimich D & Moser T (2005). Few CaV1.3 channels regulate the exocytosis of a synaptic vesicle at the hair cell ribbon synapse. J Neurosci 25, 11577–11585.

Brandt A, Striessnig J & Moser T (2003). CaV1.3 channels are essential for development and presynaptic activity of cochlear inner hair cells. J Neurosci 23, 10832–10840.

Einhorn Z, Trapani JG, Liu Q & Nicolson T (2012). Rabconnectin3α promotes stable activity of the h+ pump on synaptic vesicles in hair cells. J Neurosci 32, 11144– 11156.

Fay RR & Popper AN (2000). Evolution of hearing in vertebrates: the inner ears and processing. Hearing Research 149, 1–10.

Gale JE, Marcotti W, Kennedy HJ, Kros CJ & Richardson GP (2001). FM1-43 dye behaves as a permeant blocker of the hair-cell mechanotransducer channel. J Neurosci 21, 7013–7025.

Gallimore AR, Hepburn I, Rizzoli S & Schutter ED (2023). Dynamic regulation of vesicle pools in a detailed spatial model of the complete synaptic vesicle cycle. 2023.08.03.551909. Available at: https://www.biorxiv.org/content/10.1101/2023.08.03.551909v1 [Accessed May 1, 2024].

Giese APJ, Weng W-H, Kindt KS, Chang HHV, Montgomery JS, Ratzan EM, Beirl AJ, Rivera RA, Lotthammer JM, Walujkar S, Foster MP, Zobeiri OA, Holt JR, Riazuddin S, Cullen KE, Sotomayor M & Ahmed ZM (2023). Complexes of vertebrate TMC1/2 and CIB2/3 proteins form hair-cell mechanotransduction cation channels. eLife; DOI: 10.7554/eLife.89719.1.

Gillespie PG & Walker RG (2001). Molecular basis of mechanosensory transduction. Nature 413, 194–202.

Granato M, Eeden FJ van, Schach U, Trowe T, Brand M, Furutani-Seiki M, Haffter P, Hammerschmidt M, Heisenberg CP, Jiang YJ, Kane DA, Kelsh RN, Mullins MC, Odenthal J & Nusslein-Volhard C (1996). Genes controlling and mediating locomotion behavior of the zebrafish embryo and larva. Development 123, 399– 413.

Griesinger CB, Richards CD & Ashmore JF (2002). Fm1-43 reveals membrane recycling in adult inner hair cells of the mammalian cochlea. J Neurosci 22, 3939–3952.

Griesinger CB, Richards CD & Ashmore JF (2005). Fast vesicle replenishment allows indefatigable signalling at the first auditory synapse. Nature 435, 212–215.

Hall DH & Hedgecock EM (1991). Kinesin-related gene unc-104 is required for axonal transport of synaptic vesicles in C. elegans. Cell 65, 837–847.

Holmgren M, Ravicz ME, Hancock KE, Strelkova O, Kallogjeri D, Indzhykulian AA, Warchol ME & Sheets L (2021). Mechanical overstimulation causes acute injury and synapse loss followed by fast recovery in lateral-line neuromasts of larval zebrafish ed. Wu DK & Stainier DY. eLife 10, e69264.

Holt M, Cooke A, Neef A & Lagnado L (2004). High mobility of vesicles supports continuous exocytosis at a ribbon synapse. Curr Biol 14, 173–183.

Hummel JJA & Hoogenraad CC (2021). Specific KIF1A-adaptor interactions control selective cargo recognition. J Cell Biol 220, e202105011.

Hussain S, Pinter K, Uhl M, Wong H-T & Kindt KS (2024). Microtubule networks in zebrafish hair cells facilitate presynapse transport and fusion during development. 2024.04.12.589161. Available at: https://www.biorxiv.org/content/10.1101/2024.04.12.589161v1 [Accessed April 14, 2024].

Jing Z, Rutherford MA, Takago H, Frank T, Fejtova A, Khimich D, Moser T & Strenzke N (2013). Disruption of the presynaptic cytomatrix protein bassoon degrades ribbon anchorage, multiquantal release, and sound encoding at the hair cell afferent synapse. J Neurosci 33, 4456–4467.

Jukic A, Lei Z, Cebul ER, Pinter K, Mosqueda N, David S, Tarchini B & Kindt K (2024). Presynaptic Nrxn3 is essential for ribbon-synapse assembly in hair cells. 2024.02.14.580267. Available at: https://www.biorxiv.org/content/10.1101/2024.02.14.580267v1 [Accessed April 23, 2024].

Kamin D, Lauterbach MA, Westphal V, Keller J, Schönle A, Hell SW & Rizzoli SO (2010). High- and low-mobility stages in the synaptic vesicle cycle. Biophys J 99, 675–684.

Kimmel CB, Patterson J & Kimmel RO (1974). The development and behavioral characteristics of the startle response in the zebra fish. Dev Psychobiol 7, 47–60.

Kindt KS & Sheets L (2018). Transmission disrupted: modeling auditory synaptopathy in zebrafish. Frontiers in Cell and Developmental Biology 6, 114.

Kolla L, Kelly MC, Mann ZF, Anaya-Rocha A, Ellis K, Lemons A, Palermo AT, So KS, Mays JC, Orvis J, Burns JC, Hertzano R, Driver EC & Kelley MW (2020). Characterization of the development of the mouse cochlear epithelium at the single cell level. Nat Commun 11, 2389.

Kroll J, Jaime Tobón LM, Vogl C, Neef J, Kondratiuk I, König M, Strenzke N, Wichmann C, Milosevic I & Moser T (2019). Endophilin-A regulates presynaptic Ca2+ influx and synaptic vesicle recycling in auditory hair cells. The EMBO Journal 38, e100116.

Lee J-R et al. (2015). De novo mutations in the motor domain of kif1a cause cognitive impairment, spastic paraparesis, axonal neuropathy, and cerebellar atrophy. Human Mutation 36, 69–78.

Lenzi D, Crum J, Ellisman MH & Roberts WM (2002). Depolarization redistributes synaptic membrane and creates a gradient of vesicles on the synaptic body at a ribbon synapse. Neuron 36, 649–659.

Lenzi D, Runyeon JW, Crum J, Ellisman MH & Roberts WM (1999). Synaptic Vesicle Populations in Saccular Hair Cells Reconstructed by Electron Tomography. J Neurosci 19, 119–132.

LoGiudice L & Matthews G (2009). The role of ribbons at sensory synapses. Neuroscientist 15, 380–391.

Lukasz D & Kindt KS (2018). In vivo calcium imaging of lateral-line hair cells in larval zebrafish. J Vis Exp; DOI: 10.3791/58794.

Lush ME, Diaz DC, Koenecke N, Baek S, Boldt H, St Peter MK, Gaitan-Escudero T, Romero-Carvajal A, Busch-Nentwich EM, Perera AG, Hall KE, Peak A, Haug JS & Piotrowski T (2019). scRNA-Seq reveals distinct stem cell populations that drive hair cell regeneration after loss of Fgf and Notch signaling. Elife 8, e44431.

Lv C, Stewart WJ, Akanyeti O, Frederick C, Zhu J, Santos-Sacchi J, Sheets L, Liao JC & Zenisek D (2016). Synaptic ribbons require ribeye for electron density, proper synaptic localization, and recruitment of calcium channels. Cell Rep 15, 2784– 2795.

Marsden KC & Granato M (2015). In vivo ca(2+) imaging reveals that decreased dendritic excitability drives startle habituation. Cell Rep 13, 1733–1740.

Matthews G & Fuchs P (2010). The diverse roles of ribbon synapses in sensory neurotransmission. Nat Rev Neurosci 11, 812–822.

Meyers JR, MacDonald RB, Duggan A, Lenzi D, Standaert DG, Corwin JT & Corey DP (2003). Lighting up the Senses: FM1-43 Loading of Sensory Cells through Nonselective Ion Channels. J Neurosci 23, 4054–4065.

Michanski S, Smaluch K, Steyer AM, Chakrabarti R, Setz C, Oestreicher D, Fischer C, Möbius W, Moser T, Vogl C & Wichmann C (2019). Mapping developmental maturation of inner hair cell ribbon synapses in the apical mouse cochlea. Proceedings of the National Academy of Sciences 116, 6415–6424.

Monesson-Olson BD, Browning-Kamins J, Aziz-Bose R, Kreines F & Trapani JG (2014). Optical stimulation of zebrafish hair cells expressing channelrhodopsin-2. PLoS One 9, e96641.

Montenegro-Garreaud X et al. (2020). Phenotypic expansion in KIF1A-related dominant disorders: A description of novel variants and review of published cases. Human Mutation 41, 2094–2104.

Moser T, Brandt A & Lysakowski A (2006). Hair cell ribbon synapses. Cell Tissue Res 326, 347–359.

Mukhopadhyay M & Pangrsic T (2022). Synaptic transmission at the vestibular hair cells of amniotes. Molecular and Cellular Neuroscience 121, 103749.

Nair A, Greeny A, Rajendran R, Abdelgawad MA, Ghoneim MM, Raghavan RP, Sudevan ST, Mathew B & Kim H (2023). KIF1A-Associated Neurological Disorder: An Overview of a Rare Mutational Disease. Pharmaceuticals (Basel) 16, 147.

Neef J, Jung S, Wong AB, Reuter K, Pangrsic T, Chakrabarti R, Kügler S, Lenz C, Nouvian R, Boumil RM, Frankel WN, Wichmann C & Moser T (2014). Modes and regulation of endocytic membrane retrieval in mouse auditory hair cells. J Neurosci 34, 705–716.

Newton KC, Kacev D, Nilsson SRO, Saettele AL, Golden SA & Sheets L (2023). Lateral line ablation by ototoxic compounds results in distinct rheotaxis profiles in larval zebrafish. Commun Biol 6, 84.

Newton S, Kong F, Carlton AJ, Aguilar C, Parker A, Codner GF, Teboul L, Wells S, Brown SDM, Marcotti W & Bowl MR (2022). Neuroplastin genetically interacts with Cadherin 23 and the encoded isoform Np55 is sufficient for cochlear hair cell function and hearing. PLoS Genet 18, e1009937.

Nicolson T (2005). The genetics of hearing and balance in zebrafish. Annu Rev Genet 39, 9–22.

Niwa S, Tanaka Y & Hirokawa N (2008). KIF1Bβ- and KIF1A-mediated axonal transport of presynaptic regulator Rab3 occurs in a GTP-dependent manner through DENN/MADD. Nat Cell Biol 10, 1269–1279.

Nouvian R, Beutner D, Parsons TD & Moser T (2006). Structure and function of the hair cell ribbon synapse. J Membrane Biol 209, 153–165.

Obholzer N, Wolfson S, Trapani JG, Mo W, Nechiporuk A, Busch-Nentwich E, Seiler C, Sidi S, Söllner C, Duncan RN, Boehland A & Nicolson T (2008). Vesicular glutamate transporter 3 is required for synaptic transmission in zebrafish hair cells. J Neurosci 28, 2110–2118.

Ohta S, Ji YR, Martin D & Wu DK (2020). Emx2 regulates hair cell rearrangement but not positional identity within neuromasts. Elife 9, e60432.

Okada Y, Yamazaki H, Sekine-Aizawa Y & Hirokawa N (1995). The neuron-specific kinesin superfamily protein KIF1A is a unique monomeric motor for anterograde axonal transport of synaptic vesicle precursors. Cell 81, 769–780.

Pangrsic T & Vogl C (2018). Balancing presynaptic release and endocytic membrane retrieval at hair cell ribbon synapses. FEBS Letters 592, 3633–3650.

Paprocka J, Jezela-Stanek A, Śmigiel R, Walczak A, Mierzewska H, Kutkowska-Kaźmierczak A, Płoski R, Emich-Widera E & Steinborn B (2023). Expanding the knowledge of kif1a-dependent disorders to a group of polish patients. Genes 14, 972.

Pennings M et al. (2020). KIF1A variants are a frequent cause of autosomal dominant hereditary spastic paraplegia. Eur J Hum Genet 28, 40–49.

Petzoldt AG (2023). Presynaptic Precursor Vesicles—Cargo, Biogenesis, and Kinesin-Based Transport across Species. Cells 12, 2248.

Revelo NH, Kamin D, Truckenbrodt S, Wong AB, Reuter-Jessen K, Reisinger E, Moser T & Rizzoli SO (2014). A new probe for super-resolution imaging of membranes elucidates trafficking pathways. J Cell Biol 205, 591–606.

Rizzoli SO (2014). Synaptic vesicle recycling: steps and principles. EMBO J 33, 788– 822.

Santos MS, Li H & Voglmaier SM (2009). Synaptic vesicle protein trafficking at the glutamate synapse. Neuroscience 158, 189–203.

Schindelin J, Arganda-Carreras I, Frise E, Kaynig V, Longair M, Pietzsch T, Preibisch S, Rueden C, Saalfeld S, Schmid B, Tinevez J-Y, White DJ, Hartenstein V, Eliceiri K, Tomancak P & Cardona A (2012). Fiji: an open-source platform for biological-image analysis. Nat Methods 9, 676–682.

Schraven SP, Franz C, Rüttiger L, Löwenheim H, Lysakowski A, Stoffel W & Knipper M (2012). Altered phenotype of the vestibular organ in GLAST-1 null mice. J Assoc Res Otolaryngol 13, 323–333.

Seiler C & Nicolson T (1999). Defective calmodulin-dependent rapid apical endocytosis in zebrafish sensory hair cell mutants. Journal of Neurobiology 41, 424–434.

Sheets L, Holmgren M & Kindt KS (2021). How zebrafish can drive the future of genetic-based hearing and balance research. J Assoc Res Otolaryngol 22, 215–235.

Sheets L, Trapani JG, Mo W, Obholzer N & Nicolson T (2011). Ribeye is required for presynaptic Ca(V)1.3a channel localization and afferent innervation of sensory hair cells. Development (Cambridge, England) 138, 1309–1319.

Sidi S, Busch-Nentwich E, Friedrich R, Schoenberger U & Nicolson T (2004). gemini encodes a zebrafish L-type calcium channel that localizes at sensory hair cell ribbon synapses. J Neurosci 24, 4213–4223.

Siegel JH & Brownell WE (1986). Synaptic and Golgi membrane recycling in cochlear hair cells. J Neurocytol 15, 311–328.

Smith ET, Pacentine I, Shipman A, Hill M & Nicolson T (2020). Disruption of tmc1/2a/2b Genes in Zebrafish Reveals Subunit Requirements in Subtypes of Inner Ear Hair Cells. J Neurosci 40, 4457–4468.

Sur A, Wang Y, Capar P, Margolin G, Prochaska MK & Farrell JA (2023). Single-cell analysis of shared signatures and transcriptional diversity during zebrafish development. Dev Cell 58, 3028–3047.e12.

Trapani JG & Nicolson T (2011). Mechanism of spontaneous activity in afferent neurons of the zebrafish lateral-line organ. J Neurosci 31, 1614–1623.

Trapani JG, Obholzer N, Mo W, Brockerhoff SE & Nicolson T (2009). Synaptojanin1 is required for temporal fidelity of synaptic transmission in hair cells. PLOS Genetics 5, e1000480.

Uthaiah RC & Hudspeth AJ (2010). Molecular Anatomy of the Hair Cell’s Ribbon Synapse. J Neurosci 30, 12387–12399.

Voorn RA, Sternbach M, Jarysta A, Rankovic V, Tarchini B, Wolf F & Vogl C (2024). Slow kinesin-dependent microtubular transport facilitates ribbon synapse assembly in developing cochlear inner hair cells. 2024.04.12.589153. Available at: https://www.biorxiv.org/content/10.1101/2024.04.12.589153v1 [Accessed April 19, 2024].

Wan G, Ji L, Schrepfer T, Gong S, Wang G-P & Corfas G (2019). Synaptopathy as a mechanism for age-related vestibular dysfunction in mice. Front Aging Neurosci 11, 156.

Wichmann C & Moser T (2015). Relating structure and function of inner hair cell ribbon synapses. Cell Tissue Res 361, 95–114.

Wong HC, Zhang Q, Beirl AJ, Petralia RS, Wang Y-X & Kindt K (2019). Synaptic mitochondria regulate hair-cell synapse size and function ed. Whitfield TT, King AJ, López-Schier H & Lysakowski A. eLife 8, e48914.

Zeddies DG & Fay RR (2005). Development of the acoustically evoked behavioral response in zebrafish to pure tones. J Exp Biol 208, 1363–1372.

Zhang Q, Li S, Wong H-TC, He XJ, Beirl A, Petralia RS, Wang Y-X & Kindt KS (2018). Synaptically silent sensory hair cells in zebrafish are recruited after damage. Nat Commun 9, 1388.

Zhu S, Chen Z, Wang H & McDermott BM (2020). Tmc Reliance Is Biased by the Hair Cell Subtype and Position Within the Ear. Front Cell Dev Biol 8, 570486.

