## Supplemental Figures and Legends for "Kif1a and intact microtubules maintain synaptic-vesicle populations at ribbon synapses in zebrafish hair cells"

**Suppplemental Figures and Legends**

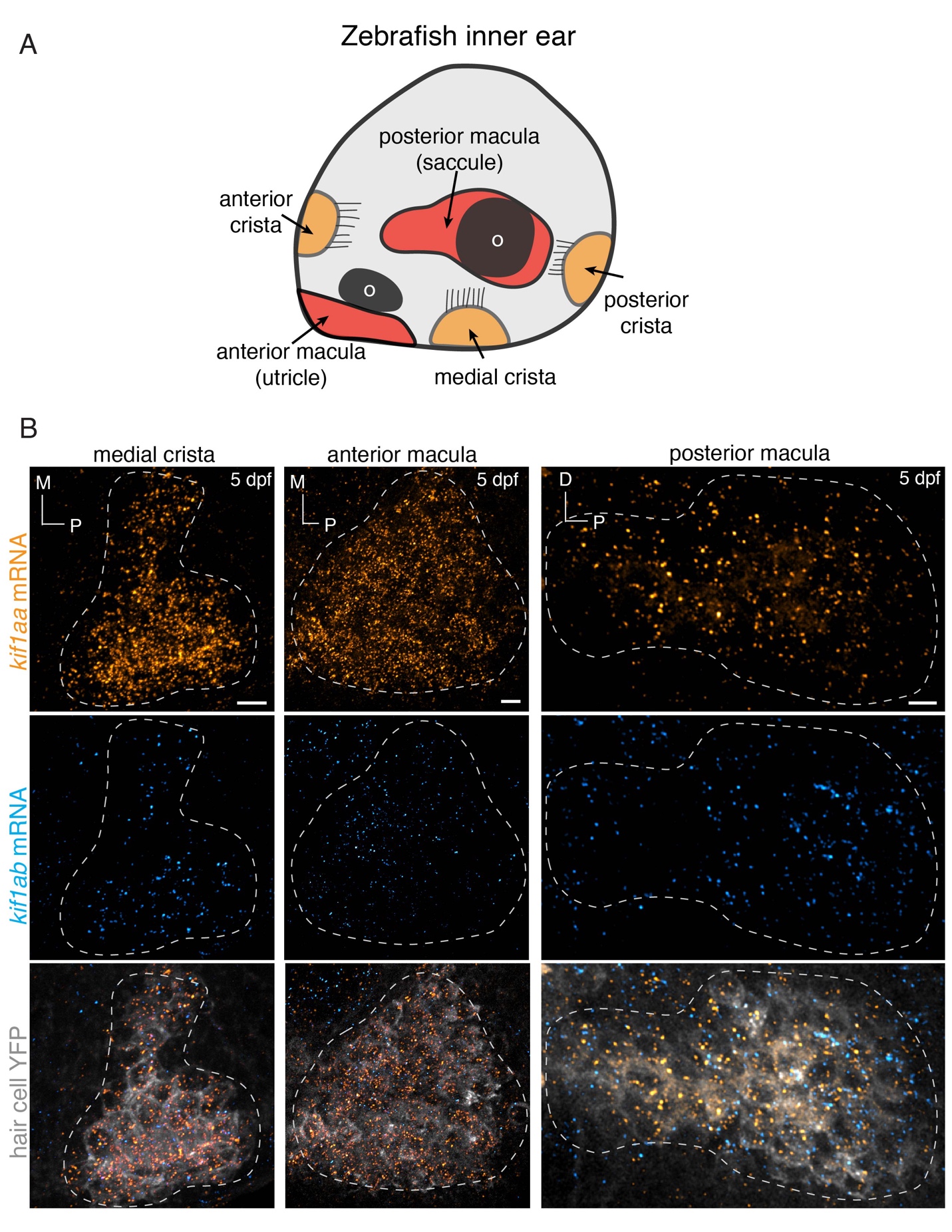

**Figure 1-S1.** ***kif1aa and kif1ab* mRNAs are both present in zebrafish inner-ear hair cells** (**A**) Schematic showing a larval zebrafish inner ear. Within the inner ear, clusters of hair cells are present in 3 cristae and 2 maculae. Each macula is associated with an otolith (o). (**B**) RNA-FISH analysis at 5 dpf reveals that both *kif1aa* (orange) and *kif1ab* (cyan) mRNAs are present in inner-ear hair cells in cristae and maculae. The gray label is YFP that is expressed specifically in hair cells. The YFP label was used to create the dashed line in **B** to outline the locations of hair cells within each sensory epithelium. Scale bars = 5 µm in **B**.

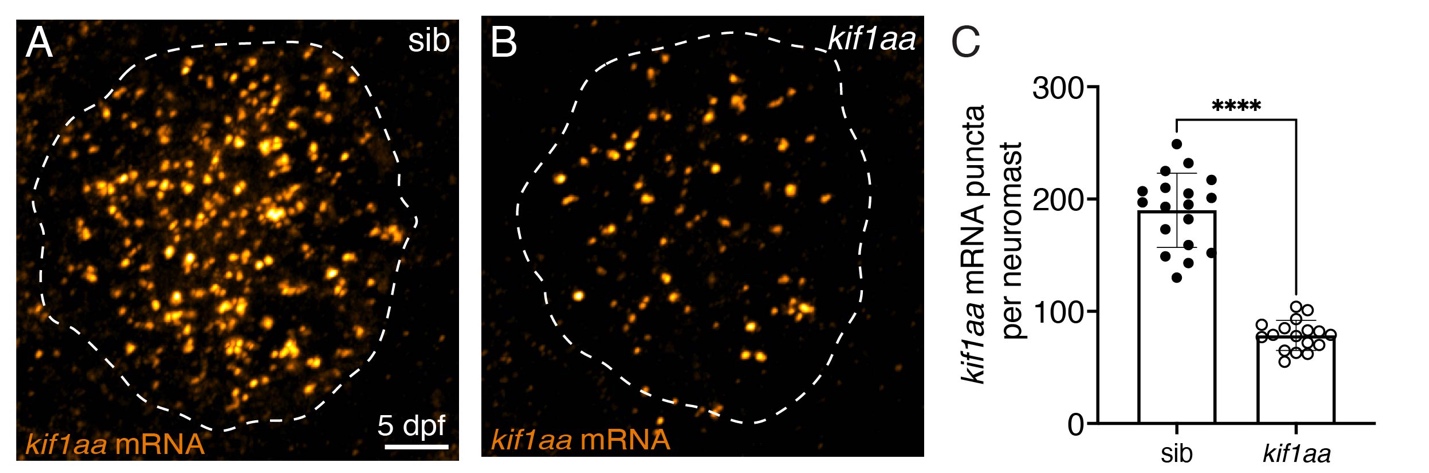

**Figure 1-S2.** ***kif1aa* mRNA levels are reduced in lateral-line hair cells in *kif1aa* mutants.**

(**A-B**) Representative RNA-FISH images of *kif1aa* mRNA in sibling control (**A**) and *kif1aa* mutant neuromasts at 5 dpf (**B**). The dashed white line indicates the neuromast boundary. (**C**) Quantification of *kif1aa* mRNA puncta reveals that the number of *kif1aa* puncta is significantly reduced in *kif1aa* mutants compared to sibling control (control: 189.9 ± 33.1; *kif1aa*: 78.6 ± 13.5, n = 18 control and 17 *kif1aa* neuromasts, unpaired t-test, p < 0.0001). Scale bar in **A** = 5 µm

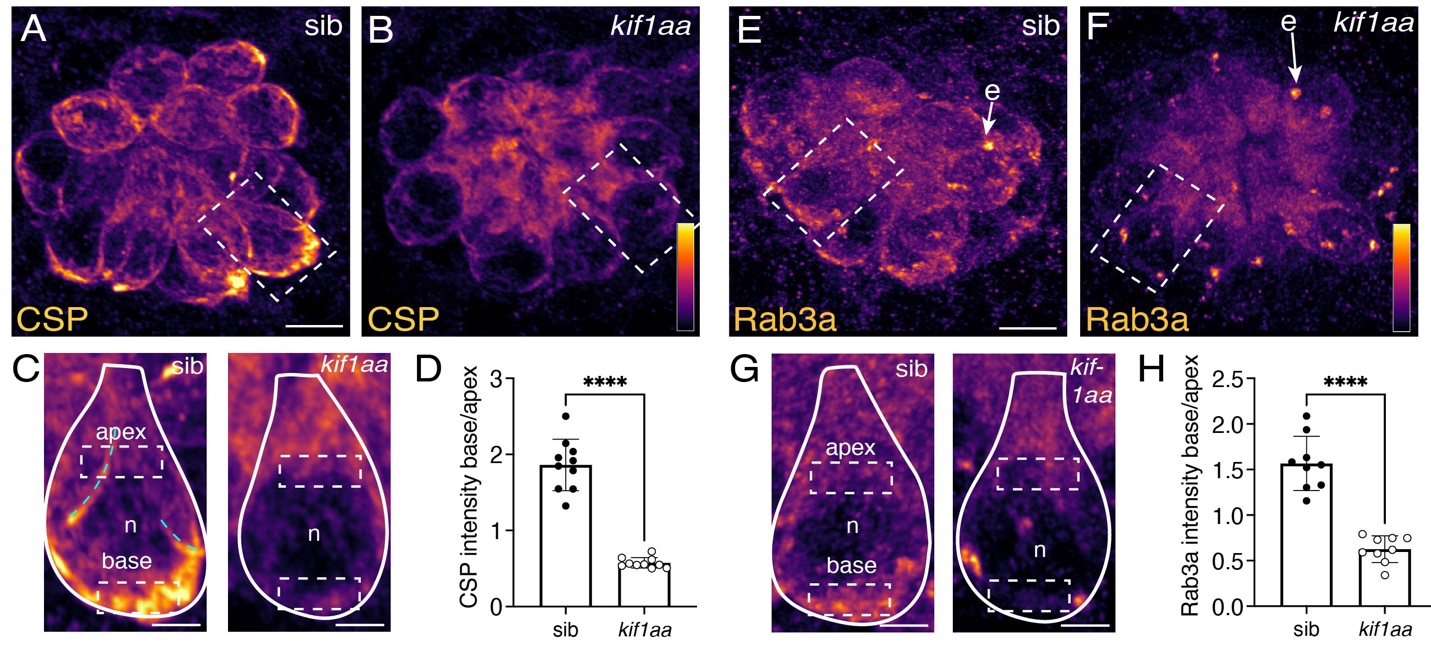

**Figure 3-S1. *Kif1aa* mutants enrich less CSP and Rab3a at the presynapse.**

(**A-C**) Example immunostain of CSP to label synaptic vesicles in neuromasts at 5 dpf in a *kif1aa* mutant (**B**) or sibling control (**A**). The dashed box in each image indicates the hair cell that is magnified and shown below in **C**. n indicates nucleus. The solid line in the magnified images outlines a single hair cell, with the base of the cell at the bottom of the image. The dashed boxes indicate example ROIs of the apical and basal regions used for intensity analysis in **D**. Cyan dashed lines indicates CSP label from neighboring cells. (**D**) Quantification of CSP label reveals that in *kif1aa* mutants, there is significantly less CSP enriched at the cell base (control: 1.86 ± 0.34, *kif1aa*: 0.58 ± 0.07, n = 10 control and *kif1aa* neuromasts, unpaired t-test, p<0.0001). (**E-G**) Example immunostain of Rab3a to label synaptic vesicles in neuromasts at 5 dpf in a *kif1aa* mutant (**F**) or sibling control (**E**). The dashed box in each images indicates the hair cell that is magnified and shown below in **G**. n indicates nucleus. e indicates efferent terminals contacting hair cells that also have high levels of Rab3a. The solid line in the magnified images outlines a single hair cell, with the base of the cell at the bottom of the image, and dashed boxes indicate example ROIs of the apical and basal regions used for intensity analysis in **H**. (**H**) Quantification reveals that the Rab3a label is significantly less enriched at the cell base (control: 1.57 ± 0.30, *kif1aa*: 0.63 ± 0.15, n = 9 control and *kif1aa* neuromasts, unpaired t-test, p<0.0001). Scale bar in **A** and **E** = 5 µm and 2 µm in **C** and **G**.

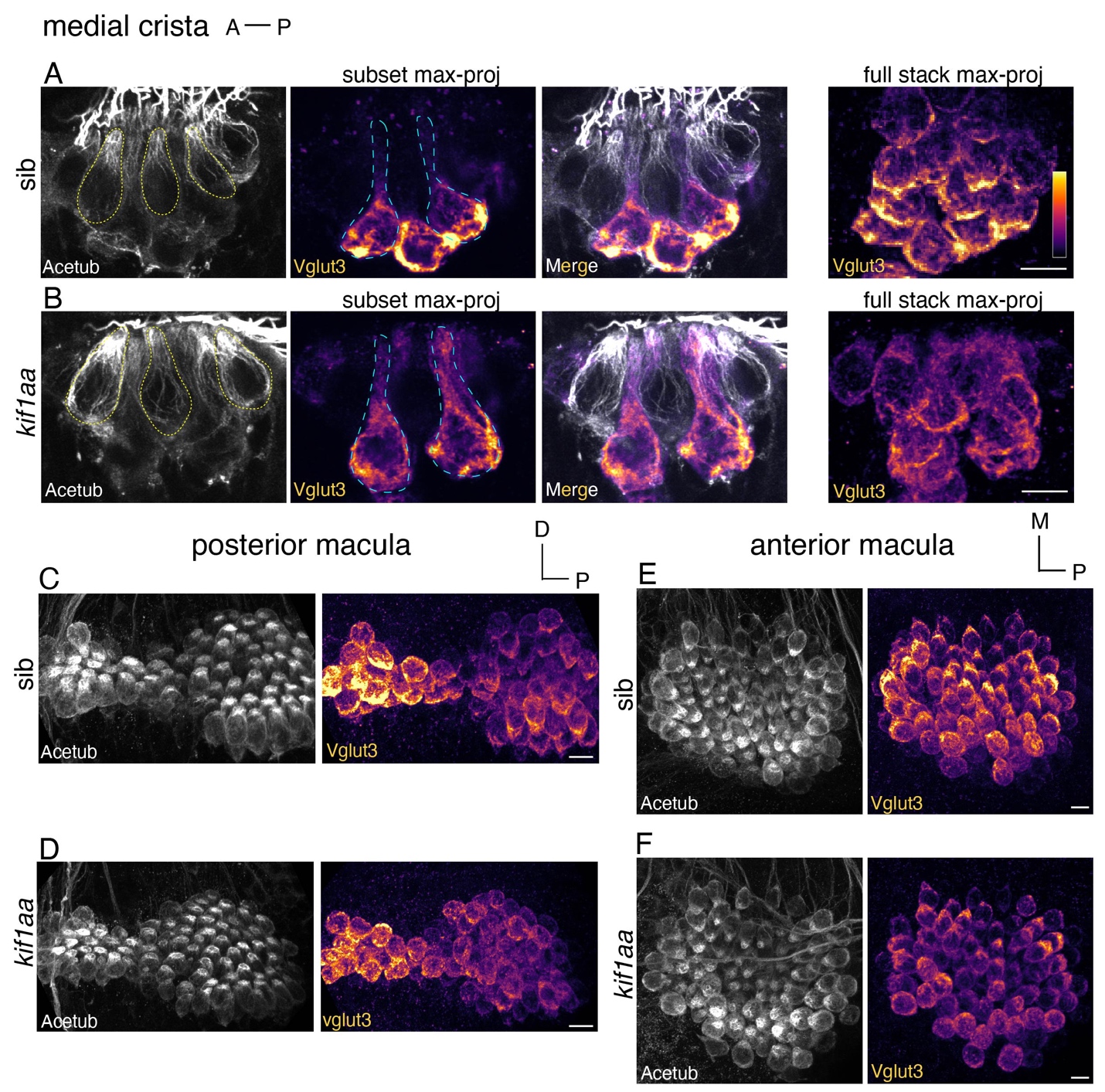

**Figure 3-S2. *Kif1aa* mutants disrupt Vglut3 localization in subsets of inner-ear hair cells.**

(**A-F**) Immunostain of inner-ear hair cells with Acetylated tubulin (Acetub, gray) to label hair cells and Vglut3 to label synaptic vesicles in the media crista (**A-B**), posterior macula (**C-D**) and anterior macula (**E-F**) in *kif1aa* mutants (**B,E,F**) and sibling controls (**A,C,D**) at 5 dpf. In the crista of both *kif1aa* mutants and controls, only a subset of hair cells (tall cells), show high levels of Vglut3 (cells outlined with cyan dashed lines in **A** and **B**). In contrast, other hair cells (tear drop cells) have no detectable Vglut3 (cells outlined with yellow dashed lines in **A** and **B**). Tall cells do not enrich Vglut3 at the cell base in *kif1aa* mutants compared to sibling controls (see partial and full stack max-projected images in **A** and **B**). In the anterior and posterior macula, Vglut3 levels are slightly reduced in *kif1aa* mutants compared to control (**C-F**). Scale bar in **A-F** = 5 µm

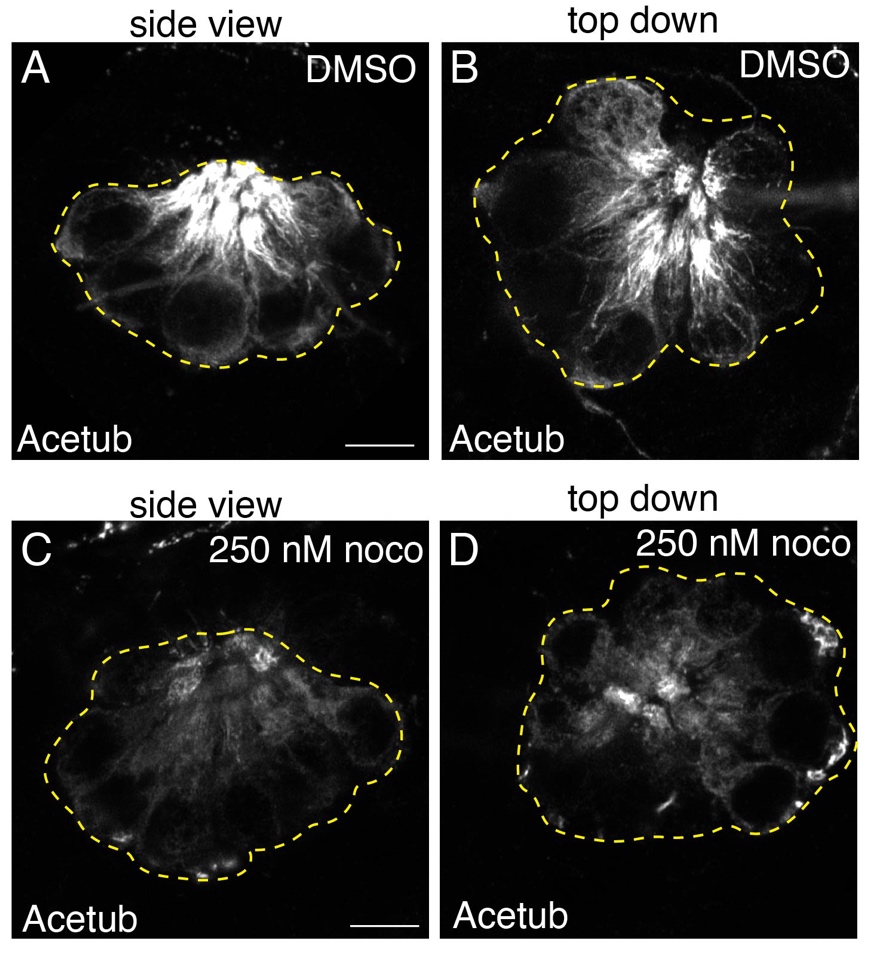

**Figure 4-S1. 250 nM nocodazole destabilizes microtubules in lateral-line hair cells.**

(**A-D**) Example fixed images of neuromasts at 5 dpf immunolabeled with Acetylated tubulin (Acetub) after a 2-hr treatment with 250 nM nocodazole (**C-D**) or after a 2 hr control treatment with 0.1% DMSO (**A-B**). Images in **A** and **C** are neuromasts viewed from the side, and **B** and **D** are viewed from the top down. The dashed lines outline the neuromasts in each image. Scale bar in **A** and **C** = 5 µm.

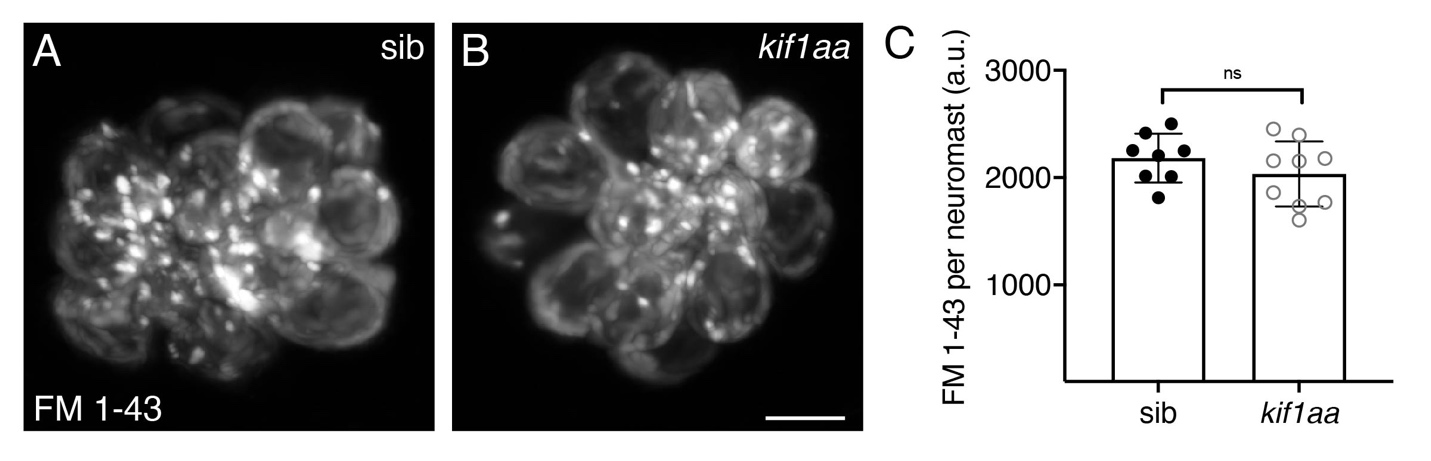

**Figure 7-S1. *kif1aa* mutant hair cells have normal FM 1-43 uptake.**

(**A-B**) Hair cells with intact mechanotransduction robustly label with the vital dye FM 1-43. Example images of FM 1-43-labeled neuromasts in sibling control (**A**) and a *kif1aa* mutant (**B**) at 5 dpf. (**C**) Quantification reveals that the average intensity of FM 1-43 label is the same between sibling controls and *kif1aa* mutants (control: 2182 ± 277, *kif1aa*: 2034 ± 302; n = 8 control and 9 *kif1aa* neuromasts, unpaired t-test, p = 0.276). Scale bar in **B** = 5 µm.

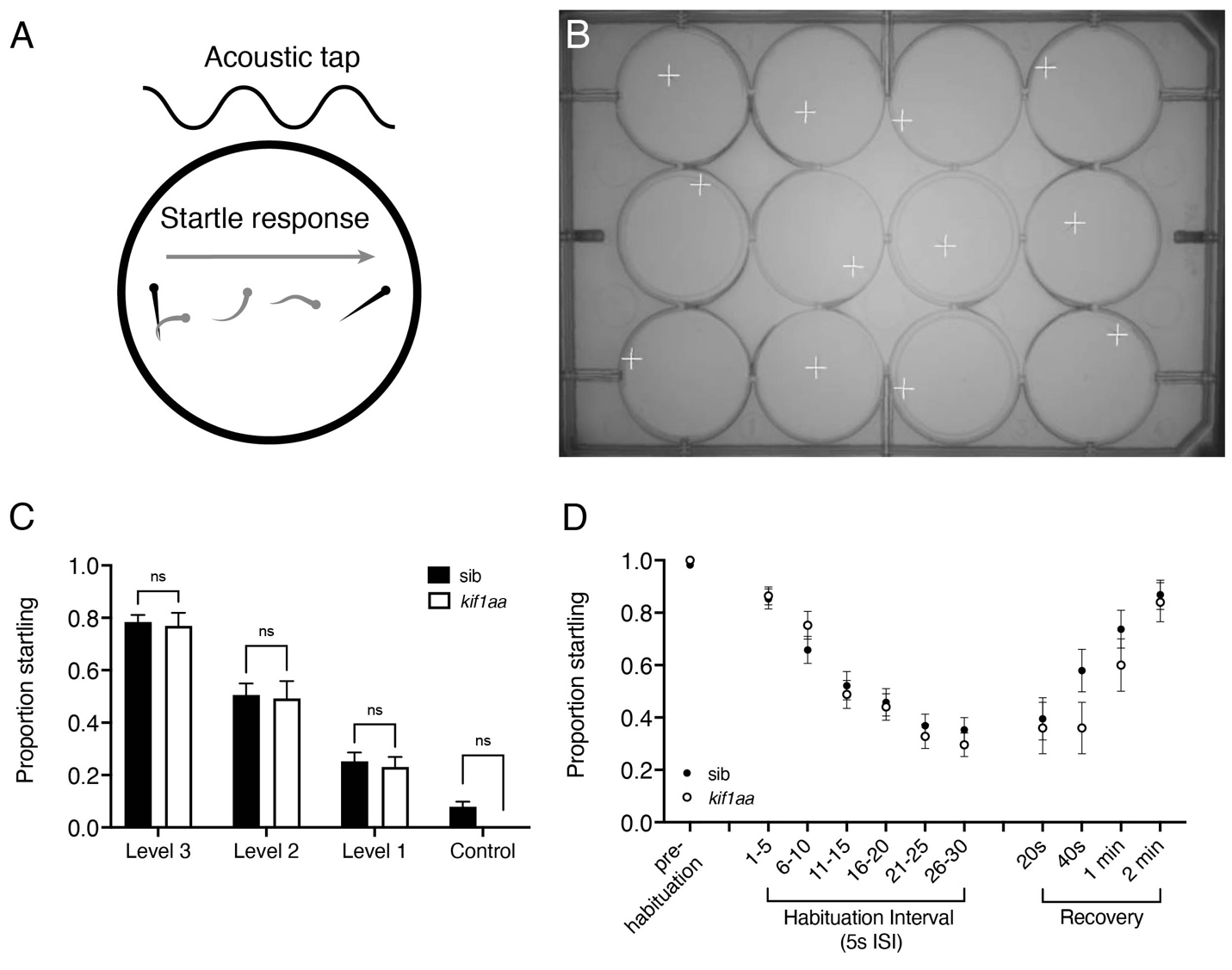

**Figure 9-S1. Acoustic startle responses are not altered in *kif1aa* mutants.**

(**A**) A vibrational acoustic tap stimulus was used to evoke an escape response in sibling control and *kif1aa* mutant larvae in each well of a 12-well plate. (**B**) In each plate, the 12 fish were tracked using Zantiks software, and the response to each stimulus was recorded. (**C**) A vibrational tap of 3 levels of decreasing intensity, with 5 stimuli per level of intensity, was administered and the proportion of times each animal responded was plotted. There is no significant difference between *kif1aa* mutants and sibling controls in responding to a single tap stimulus at any intensity tested (control: 0.78 ± 0.16 (Level 3), 0.51 ± 0.27 (Level 2), 0.25 ± 0.21 (Level 1), 0.08 ± 0.12 (no stimulus); *kif1aa*: 0.77 ± 0.18 (Level 3), 0.49 ± 0.24 (Level 2), 0.23 ± 0.14 (Level 1), 0 ± 0 (no stimulus); n = 38 control and 13 *kif1aa* larvae, 2-way ANOVA with multiple comparisons, p = 0.999 (Level 3-2), 0.994 (Level 1), and p = 0.586 for no stimulus control; 5 dpf). (**D**) In a habituation assay *kif1aa* mutants do not habituate or recover after habituation at a significantly different rate compared to sibling controls (n = 38 control and 25 *kif1aa* larvae, 2-way ANOVA with multiple comparisons, habituation: genotype x stimulus, p = 0.545 (and no significant difference at any interval); recovery: 40 s, p = 0.318; 1 min, p = 0.721; 2 min, p = 0.997).

| GLMM | Estimated | Std. Error | df | t value | Pr(>\|t\|) |
| --- | --- | --- | --- | --- | --- |
| (Intercept) | 8.365 | 3.51 | 209.881 | 2.383 | **1.81E-02*** |
| Stimulus 10s: Genotype mutant -/- | -9.249 | 7.463 | 141.067 | -1.239 | 0.2173 |
| Stimulus 20s: Genotype mutant -/- | -18.121 | 7.503 | 141.945 | -2.415 | **1.70E-02*** |

| ANOVA (III) | Sum Sq | Mean Sq | NumDF | DenDF | F value | Pr(>F) |
| --- | --- | --- | --- | --- | --- | --- |
| Stimulus | 46726 | 23363 | 2 | 141.65 | 47.48 | < 2e-16*** |
| Genotype | 1510 | 1509.8 | 1 | 71.079 | 3.0684 | 0.08414 |
| Stimulus:Genotype | 2871 | 1435.7 | 2 | 141.65 | 2.9177 | 0.05731 |

**Supplementary Table 1**. Generalized Linear Mixed Model with Satterthwaite’s method of post hoc *t-*tests for differences in the total distance traveled during rheotaxis events. Type III ANOVA yielded significance values for fixed effects because the LME4 package in R does not identify them in its output. Significance codes: ‘***’ 0.001, ‘*’ 0.05

| GLMM | Estimated | Std. Error | df | t value | Pr(>\|t\|) |
| --- | --- | --- | --- | --- | --- |
| (Intercept) | 0.09302 | 0.27339 | 204.5077 | 0.34 | 0.734 |
| Stimulus 10s: Genotype mutant -/- | -0.34961 | 0.55798 | 142 | -0.627 | 0.5319 |
| Stimulus 20s: Genotype mutant -/- | -1.00853 | 0.55798 | 142 | -1.807 | 0.0728 |

| ANOVA (III) | Sum Sq | Mean Sq | NumDF | DenDF | F value | Pr(>F) |
| --- | --- | --- | --- | --- | --- | --- |
| Stimulus | 592.56 | 296.278 | 2 | 142 | 107.704 | <2e-16*** |
| Genotype | 2.74 | 2.743 | 1 | 71 | 0.9972 | 0.3214 |
| Stimulus:Genotype | 9.27 | 4.634 | 2 | 142 | 0.1892 | 0.1892 |

**Supplementary Table 2**. Generalized Linear Mixed Model with Satterthwaite’s method of post hoc *t-*tests for differences in the mean number of rheotaxis events. Type III ANOVA yielded significance values for fixed effects because the LME4 package in R does not identify them in its output. Significance codes: ‘***’ 0.001, ‘*’ 0.05
